# Bursts of coalescence within population pedigrees whenever big families occur

**DOI:** 10.1101/2023.10.17.562743

**Authors:** Dimitrios Diamantidis, Wai-Tong (Louis) Fan, Matthias Birkner, John Wakeley

## Abstract

We consider a simple diploid population-genetic model with potentially high variability of offspring numbers among individuals. Specifically, against a backdrop of Wright-Fisher reproduction and no selection there is an additional probability that a big family occurs, meaning that a pair of individuals has a number of offspring on the order of the population size. We study how the pedigree of the population generated under this model affects the ancestral genetic process of a sample of size two at a single autosomal locus without recombination. Our population model is of the type for which multiple-mergers coalescent processes have been described. We prove that the conditional distribution of the pairwise coalescence time given the random pedigree converges to a limit law as the population size tends to infinity. This limit law may or may not be the usual exponential distribution of the Kingman coalescent, depending on the frequency of big families. But because it includes the number and times of big families it differs from the usual multiple-merger coalescent models. The usual multiple-merger coalescent models are seen as describing the ancestral process marginal to, or averaging over, the pedigree. In the limiting ancestral process conditional on the pedigree, the intervals between big families can be modeled using the Kingman coalescent but each big family causes a discrete jump in the probability of coalescence. Analogous results should hold for larger samples and other population models. We illustrate these results with simulations and additional analysis, highlighting their implications for inference and understanding of multi-locus data.

## Introduction

### Population-genetic background

Population geneticists routinely make inferences about the past by applying statistical models to DNA sequences or other genetic data. Because past events have already occurred, these models describe what might have happened. They are necessary because patterns of variation in DNA provide only indirect evidence about the past. But the decisions made in building these statistical models have important consequences for inference. A key question has received little attention: when and how should some parts of the past be treated as random variables, while others are viewed as fixed objects? Our particular concern here will be with the treatment of pedigrees, or the reproductive relationships among diploid individuals.

With limited exceptions the statistical models of population genetics have inherited the initial decisions which Fisher (1922, 1930) and Wright (1931) made in deriving allele frequency spectra and probability density functions of allele frequencies at stationarity. They modeled neutral alleles as well as those under selection in a large well-mixed population which in the simplest case was assumed to be of constant size over time. Accordingly it has been common in population genetics to think of population sizes as fixed, not random. Today’s coalescent hidden Markov models, for example, infer a fixed trajectory of population sizes over time under the assumption of neutrality (Li and Durbin, 2011; Sheehan et al., 2013; Wang et al., 2020; Schweiger and Durbin, 2023).

Although coalescent models reflect later developments and were a significant shift in thinking for the field, fundamentally they depend on the same assumptions as the classical models of Fisher and Wright (Ewens, 1990; Möhle, 1999). This is clear even in the earliest treatments of ancestral genetic processes by Malécot (1941, 1946, 1948). What coalescent theory did was to broaden the scope of population genetics beyond forward-time models of changes in allele counts or frequencies to include gene genealogies constructed by series of common-ancestor events backward in time (Kingman, 1982; Hudson, 1983a,b; Tajima, 1983). Mathematically, the forward-time and backward-time models of population genetics are dual to each other (Möhle, 1999).

Most importantly for our purposes here, Fisher (1922, 1930) and Wright (1931) obtained their predictions about genetic variation by averaging over an assumed random process of reproduction. The particular random process they used is now called the Wright-Fisher model (Ewens, 2004). Because the outcome of the process of reproduction is a pedigree, their method is equivalent to averaging over the random pedigree of the population. That they did this without explanation in this context is somewhat curious given the attention to pedigrees in Fisher’s infinitesimal model of quantitative genetics (Fisher, 1918; Barton et al., 2017) and in Wright’s method of path coefficients whose very purpose was to make predictions conditional on pedigrees (Wright, 1921a,b,c,d,e, 1922).

The pedigree of the entire population is the set of reproductive relationships of all individuals for all time when reproduction is bi-parental. The corresponding graph is a genealogy in the usual sense. It has been referred to as an organismal pedigree (Ball et al., 1990) and the population pedigree (Wollenberg and Avise, 1998; Wakeley et al., 2012; Ralph, 2019). Here we simply call it the pedigree. Patterns of genetic variation depend on the pedigree because genetic inheritance happens within it. In particular, transmission of an autosomal genetic locus forward in time through the pedigree occurs by Mendel’s law of independent segregation. Multi-locus transmission follows Mendel’s law of independent assortment or is mediated by recombination if the loci are linked. These processes, which may also be viewed backward in time, are conditional on the pedigree.

Within any pedigree, many possible uni-parental paths can be traced backward in time from each individual. If there are two mating types, for example karyotypic females (F) and karyotypic males (M), then one such path might be depicted F→F→M→F→M→ *· · ·* (Avise and Wollenberg, 1997). For the ancestry of a single allele at an autosomal locus in a single individual, applying Mendel’s law of independent segregation backward in time generates these uni-parental paths with equal probabilities 1/2^*g*^ for any path extending *g* generations into the past. When two such paths meet in the same individual, then with equal probability, 1/2, the alleles either coalesce in that individual or remain distinct. Thus coalescence is conditional on the pedigree, and many possible gene genealogies are embedded in any one pedigree. Some loci, such as the mitochondrial genome and the Y chromosome in humans, are strictly uni-parentally inherited. They follow only paths F→F→F→ *· · ·* and M→M→M→ *· · ·*, respectively, and two such paths coalesce with probability one when they meet. For these loci, there is only one gene genealogy within the pedigree.

Under Wright-Fisher reproduction, parents are chosen at random uniformly from among all possible parents. This determines the structure of the pedigree in that generation. Assume that there are *N*_f_ karyotypic females and *N*_m_ karyotypic males in every generation. For autosomal loci, the familiar effective population size *N*_*e*_ = 4*N*_f_ *N*_m_/(*N*_f_ + *N*_m_) from classical forward-time analysis (Wright, 1931) and its backward-time counterpart 1/(2*N*_*e*_) for the pairwise coalescence probability (Möhle, 1998a,b) come from averaging over the possible outcomes of reproduction in a single generation. Sections 6.1 and 6.2 in Wakeley (2009) give a detailed illustration. For uni-parentally inherited loci, this averaging yields 1/*N*_f_ and 1/*N*_m_ for the pairwise coalescence probabilities. In the diploid monoecious Wright-Fisher model or by setting *N*_f_ = *N*_m_ = *N*/2, these average probabilities of coalescence become 2/*N* for uni-parentally inherited loci and 1/(2*N*) for autosomal loci. For simplicity in this work we will focus on the diploid monoecious Wright-Fisher model.

Averaging over pedigrees is what leads to the effective population size, *N*_*e*_, being the primary determinant of forward-time and backward-time dynamics in neutral population genetic models. For very large populations, *N*_*e*_ becomes the only parameter of the Wright-Fisher diffusion (Ewens, 2004) and the standard neutral or Kingman coalescent process (Sjödin et al., 2005). In particular, *N*_*e*_ sets the timescale over which mutation acts to produce genetic variation. Such averaging removes the pedigree as a possible latent variable which could be important for structuring genetic variation. As a result, from the perspective of the standard neutral coalescent, information about the (marginal) gene genealogical process together with the mutation process is all we can hope to infer from genetic data (Sjödin et al., 2005).

The situation in which it makes the most sense to use this marginal process of coalescence is when the only data available come from a single non-recombining locus. In fact, the initial applications of ancestral inference to single-locus data, namely to restriction fragment length polymorphisms in human mitochondrial DNA (mtDNA) (Brown, 1980; Cann et al., 1987) then to sequences of the hyper-variable control region (Vigilant et al., 1989, 1991; Ward et al., 1991; Di Rienzo and Wilson, 1991), did not even use of the statistical machinery of population genetics. They instead took the gene genealogy and times to common ancestry to be fixed, and estimated them using traditional phylogenetic methods (Felsenstein, 2004). But this in turn spurred the development of likelihood-based methods of ancestral inference using coalescent prior distributions for gene genealogies (Lundstrom et al., 1992; Griffiths and Tavaré, 1994; Kuhner et al., 1995). We note that in the interim it has also become common to treat phylogenies as random variables using a wide variety of prior models (Ronquist et al., 2012; Suchard et al., 2018; Bouckaert et al., 2019).

The desirability of accounting for variation in gene genealogies became especially clear when the first sample DNA sequences of the human *ZFY* gene was obtained and was completely monomorphic (Dorit et al., 1995). The mutation rate is lower on the Y chromosome than in the hyper-variable region of mtDNA but it is not equal to zero (Brown et al., 1979; Wilson et al., 1985; Ingman et al., 2000; The 1000 Genomes Project Consortium, 2015). Using coalescent priors it was shown that the complete lack of variation in that first sample at *ZFY* was consistent with a wide range of times to common ancestry for the Y chromosome (Dorit et al., 1995; Donnelly et al., 1996; Fu and Li, 1996; Weiss and von Haeseler, 1996).

If instead data come from multiple loci, it is impossible to ignore variation in gene genealogies regardless of whether one thinks of the pedigree as fixed or random. Variation in gene genealogies across the genome is, for example, what coalescent hidden Markov models use to estimate trajectories of population sizes. The simplest illustrative case is when the loci are on different chromosomes or far enough apart on the same chromosome that they assort independently into gametes, and when within each locus there is no recombination. The gene genealogies of such loci will vary due to the particular outcomes of Mendelian segregation. They will also be independent due to Mendelian assortment, but only given the pedigree. Mendel’s law of independent assortment is a law of conditional independence. It applies once relationships have been specified.

However, throughout much of the history of population genetics, it was assumed that independently assorting loci would have completely independent evolutionary histories. In coalescent theory, this means independent gene genealogies. As Charlesworth (2022) recently noted, Fisher (1922, 1930) and Wright (1931) intended their results on allele frequency spectra and probability density functions of allele frequencies at stationarity to be descriptions of the behavior of large numbers of independently assorting loci in the same genome. This is evident in their application of these distributions to the multiple Mendelian factors of Fisher’s infinitesimal model (Fisher, 1918) in their arguments about the Dominance Ratio (Fisher, 1922; Charlesworth, 2022).

An early application to multi-locus data was made by Cavalli-Sforza and Edwards (1967) and Felsenstein (1973) who developed likelihood-based methods to infer trees of populations within species from multi-locus allele-frequency data, specifically human blood group data, by modeling the forward-time process of random genetic drift independently at each locus conditional on the population tree. Felsenstein (1981) further developed and applied these methods to gel electrophoretic data. Today’s methods of inferring admixture from single nucleotide polymorphism, or SNP, data using F-statistics are based on the same notion of independence (Patterson et al., 2012).

Like the population size itself, demographic features such as the splitting of populations have mostly been treated as fixed in population genetics. Cavalli-Sforza and Edwards (1967) and Felsenstein (1973) did discuss but did not implement prior models for trees of populations, specifically as outcomes of birth-death processes. More recently, Heled and Drummond (2009) did implement this in a coalescent framework for multi-locus sequence data, using the prior distribution of Gernhard (2008); see also Lambert and Stadler (2013). Yang (2002) and Rannala and Yang (2003) took a different approach, using gamma-distributed pseudo priors for times in trees.

### Previous work on pedigrees

Although the underlying assumption that unlinked loci have completely independent evolutionary histories is mistaken because it would require them having independent pedigrees, most theoretical work has followed the lead of Fisher (1922, 1930) and Wright (1931). Examples in which this is made explicit include Karlin and McGregor (1967), Kimura (1969), Ewens (1974), and Ewens and Maruyama (1975). Multiplying likelihoods across loci in applications to genetic data subsequently became common practice (Watterson, 1985; Pad-madisastra, 1988; Sawyer and Hartl, 1992; Wakeley, 1999; Nielsen, 2000; Wooding and Rogers, 2002; Adams and Hudson, 2004). It is built into current inference packages, including *∂*a*∂i* (Gutenkunst et al., 2009), *momi* 2 (Kamm et al., 2020) and *fastsimcoal* 2 (Excoffier et al., 2013, 2021).

As it happens, this conceptual mistake has almost no practical ramifications if the population is large and well mixed, and the variance of offspring numbers among individuals is not too large. Ball et al. (1990) were the first to address the question of gene genealogies within pedigrees. They used simulations to show that the distribution of pairwise coalescence times among loci on a single pedigree do not differ substantially from their distributions among loci which have independent pedigrees. Their population model was similar to the Wright-Fisher model with population size *N* = 100: Poisson offspring numbers with strong density regulation to a carrying capacity of 100. Their results were based on simulations of 50 gene genealogies for each of 50 pedigrees and samples of size *n* = 100, in which a single gene copy was taken at random at each locus within each individual. They also showed that the distribution of coalescence times among pairs of individuals on a single pedigree are very similar to the prediction obtained by averaging over pedigrees.

Wakeley et al. (2012) confirmed these results and related them to coalescent theory using simulations of 10^8^ gene genealogies for *n* = 2 for each of 10^4^ pedigrees and population sizes up to 10^5^, together with more limited treatments of larger samples *n* = 20 and *n* = 100. Pedigrees were constructed in three different ways: assuming Wright-Fisher reproduction, using empirically derived human family structures, and under a model in which the outcome of a single generation of Wright-Fisher reproduction was repeated over time, resulting in a so-called cyclical pedigree. These simulations showed that times to common ancestry conditional on the pedigree conform well to the probability law underlying coalescent theory, with a constant coalescence probability 1/(2*N*_*e*_) = 1/(2*N*) each generation under the Wright-Fisher model with *N*_f_ = *N*_m_ = *N*/2, except for in the recent past where they differ greatly and depend on the pedigree. But they also showed that as long as *N* is large these idiosyncrasies in the short-time behavior of the ancestral process have little effect on the overall distribution of coalescence times given the pedigree, whether it is among independent loci in the same individuals or among independently sampled pairs of individuals.

Here “recent” means proportional to log_2_(*N*) generations, which is the timescale for the first occurrence of a common ancestor of all present-day individuals (Chang, 1999) and for the complete overlap of all individuals’ ancestries in a well-mixed bi-parental population (Chang, 1999; Derrida et al., 1999, 2000a,b; Barton and Etheridge, 2011; Coron and Le Jan, 2022). This is much shorter than the *N*-generations timescale required for common ancestry of uni-parental genetic lineages (Chang, 1999; Donnelly et al., 1999). Further work on these properties of pedigrees include Rohde et al. (2004) and Lachance (2009) who showed that population structure and inbreeding do not strongly affect the time to the first occurrence of a common ancestor of all individuals. Blath et al. (2014) proved that the ancestries of the great majority of individuals overlap even in cyclical pedigrees as *N* → *∞*. Matsen and Evans (2008) and Gravel and Steel (2015) showed that ancestral genetic lineages pass through only a small minority of the shared pedigree ancestors. Sainudiin et al. (2016) constructed a model with recombination which interpolates between uni-parental common ancestry on the *N*-generations timescale and bi-parental common ancestry on the log_2_(*N*)-generations timescale.

Tyukin (2015) proved what was implied by the simulations of Ball et al. (1990) and Wakeley et al. (2012), specifically that when the population is large and well mixed the pedigree-averaged coalescent process is a good substitute for the actual coalescent process conditional on the pedigree. Questions of this sort have a long history in mathematical physics and probability theory, where “quenched” and “annealed” are often used to refer to conditional as opposed to averaged processes. Molchanov (1994) and Bolthausen and Sznitman (2002b) provide background and developments in the classical context of random walks in random environments. What Tyukin (2015) proved is that the quenched coalescent process conditional on the pedigree converges to the pedigree-averaged standard neutral or Kingman coalescent process in the limit *N* → *∞*. Tyukin (2015) did this under a broader set of reproduction models with mating analogous to Wright-Fisher but with a general exchangeable distribution of offspring numbers (Cannings, 1974) in the domain of attraction of Kingman’s coalescent (Möhle and Sagitov, 2001; Sagitov, 2003).

Since time in the Kingman coalescent process is measured in units proportional to *N* generations, the result of Tyukin (2015) provides insight into the role of the pedigree in the recent ancestry of the sample (*∝* log_2_(*N*) generations) under the Cannings and Wright-Fisher models. Specifically, the chance of any events in the recent past which would dramatically alter the rate of coalescence must be negligible as *N* → *∞*. Intuitively we might surmise that (1) individuals randomly sampled from a large well mixed population are unlikely to be closely related, and barring coalescence for some small number of generations until their ancestries overlap does not affect the limit, and (2) by the time their ancestries do overlap in the pedigree, their numbers of ancestors are approaching the population size, making the chance of coalescence of order 1/*N*.

Two cases have been identified using simulations where quenched and annealed results are noticeably different. The first is population subdivision, especially with limited migration. Wollenberg and Avise (1998) showed that as the migration distance decreases in a linear habitat, fewer independent loci are needed to accurately measure pairwise coefficients of coancestry on the pedigree. Wilton et al. (2017) described increasingly strong pedigree effects as the migration rate decreased in a two-subpopulation model, specifically spikes in the distribution of pairwise coalescence times corresponding to the particular series of individual migration events that occurred in the ancestry. These results illustrate how even single gene genealogies may contain information about events in the ancestry of geographically structured populations, via the pedigree. Thus they are relevant for applications of ancestral inference to single-locus data, such as mtDNA, as well as to the broader field of intraspecific phylogeography (Avise et al., 1987; Avise, 1989, 2000). For recent empirical studies of spatiotemporally structured pedigrees and their effects on local patterns of genetic variation, see Aguillon et al. (2017) and Anderson-Trocmé et al. (2023).

The second situation in which pedigrees have a strong effect on coalescence times and gene genealogies is when there is a high variance of offspring numbers among individuals. This variance is comparatively low in the Wright-Fisher model, which has a multinomial distribution of offspring numbers (becoming Poisson as *N* → *∞*). In deriving the standard neutral coalescent process, Kingman (1982) started with the general exchangeable model of Cannings (1974) then assumed that the variance of offspring numbers was finite as *N* → *∞*. Without this assumption, the ancestral limit process is not the Kingman coalescent process but rather a coalescent process with multiple mergers (see below). In addition in this situation simulations have shown that the pedigree has a marked effect on genetic ancestries.

Wakeley et al. (2016) simulated pedigrees in which a single individual had a very large number of offspring in some past generation and otherwise there was Wright-Fisher reproduction. This large reproduction event greatly increased the probability of coalescence in the generation in which it occurred, causing a spike in the distribution of pairwise coalescence times and altering the allele frequency spectrum. A strong selective sweep at one locus gave similar effects at unlinked loci via the pedigree (Wakeley et al., 2016). Similar deviations from standard neutral coalescent predictions are produced by cultural transmission of reproductive success (Guez et al., 2023).

### Plan of the present work

Here, we present a new quenched limit result for coalescent processes in fixed pedigrees under a modified Wright-Fisher model which allows for large reproduction events. Wright-Fisher reproduction on its own produces various kinds of large reproduction events but these are all extremely rare. Our model adds big families with two parents and numbers of offspring proportional to the population size. These are inserted into the pedigree either on the same *N*-generations timescale as coalescent events in the Wright-Fisher background model or much faster so that they completely dominate the ancestral process. In both cases, the limiting ancestral process conditional on the pedigree is different than the limiting ancestral process which averages over pedigrees. For simplicity, we focus on samples of size two. Consistent with the results of Tyukin (2015), our result reduces to the Kingman coalescent with *n* = 2 in the case where there are no big families.

Note that the corresponding averaged process is not the Kingman coalescent but rather a coalescent process with multiple mergers; see Tellier and Lemaire (2014) for an overview of these models in the context of population genetics. Multiple-mergers coalescent processes arise as *N* → *∞* limits when the variance of offspring numbers is large, and so may be applicable to a broad range of species with the capacity for high fecundity (Eldon, 2020). They also arise from recurrent selective sweeps, when differences in offspring numbers are determined by individuals’ genotypes (Durrett and Schweinsberg, 2004, 2005; Schweinsberg and Durrett, 2005). Whereas the Kingman coalescent includes only binary mergers of ancestral genetic lineages, these more general processes allow mergers of any size. At issue here is how these models should be interpreted and applied.

By averaging over the process of reproduction, two kinds of multiple-mergers coalescent processes have been described: Λ-coalescents which have asynchronous multiple-mergers (Donnelly and Kurtz, 1999; Pitman, 1999; Sagitov, 1999) and Ξ-coalescents which have simultaneous multiple-mergers (Schweinsberg, 2000; Möhle and Sagitov, 2001; Sagitov, 2003). Multiple-mergers processes for diploid organisms are always Ξ-coalescents with the possibility of an even number simultaneous mergers (Birkner et al., 2018). Our quenched limit result brings into question what seems like a natural extension from applications of the standard neutral coalescent model, namely to assume that multiple-mergers models may be applied independently to independent loci as has been done both in theoretical explorations (Der and Plotkin, 2014; Eldon et al., 2015; Spence et al., 2016; Matuszewski et al., 2018) and in analyses of SNP data (Birkner et al., 2013a; Blath et al., 2016; Árnason et al., 2023; Freund et al., 2023).

To establish the quenched limit process, we adapt the method that Birkner et al. (2013c) used for a quenched limit of a random walk in a random environment. See also the earlier work of Bolthausen and Sznitman (2002a). In this approach, the problem of convergence in distribution is addressed by analyzing a pair of conditionally independent processes, here corresponding to the ancestries of samples at two independently assorting loci on the pedigree. As Koskela (2018) has pointed out, positive correlations of coalescence times for pairs of unlinked loci are a hallmark of (pedigree-averaged) multiple-mergers coalescent models. Our result frames this in terms of pedigrees, in which big families are the only elements that persist as *N* → *∞*. If a big family has occurred in a particular generation, the probability of coalescence is greatly increased in that generation for all loci. All other aspects of the pedigree, that is to say the outcomes of ordinary Wright-Fisher reproduction, “average out” such that the Kingman coalescent process describes the ancestral process during the times between big families.

### Theory and results

In this section, we present the population model considered in this paper, the mathematical statement of our main result and its proof. This result (Theorem 1) is stated as a convergence of the *conditional* distribution of the coalescence time of a pair of gene copies, given that we know the pedigree and which individuals were sampled. We assume that the pedigree is the outcome of a random process of reproduction, the population model described in the following section, and that the two individuals are sampled without replacement from the current generation. To connect with known results and highlight the effect of conditioning, we first state and prove the corresponding result (Lemma 1) for the *un*conditional distribution of the coalescence time. This corresponds to fixing the sampled individuals and averaging over the pedigree. We close this section with simulations illustrating multi-locus genetic ancestry and further analysis showing how non-zero correlations of coalescence times at unlinked loci result from averaging over the pedigree.

### The population model

We consider a diploid, monoecious, bi-parental, panmictic population of constant fixed size *N* ∈ N with discrete, non-overlapping generations. Implicitly there is no selection, but we do not in fact model mutation or genetic variation, only the generation of the pedigree and coalescence within it. There are two different types of reproduction. With high probability, reproduction follows the diploid bi-parental Wright-Fisher model. With small probability *α*_*N*_ each generation, there is a highly reproductive pair whose offspring comprise a proportion *ψ* ∈ [0, 1] of the population. Note that *ψ* is a fixed deterministic constant. More precisely, for each positive integer *g*, the reproductive dynamics between the parent generation *g* + 1 and the offspring generation *g* is given as follows:

1. With probability 1 − *α*_*N*_, each individual in the next generation is formed by choosing two parents at random, uniformly with replacement from the *N* adults of the current generation. Genetically, each offspring is produced according to Mendel’s laws which means each of the two gene copies in a parent is equally likely to be the one transmitted to the offspring. In this case we call *g* a *“Wright-Fisher generation”*. An example of this standard reproduction dynamics between the parent generation *g*+1 and the offspring generation *g* is depicted below for a population of size *N* = 7.

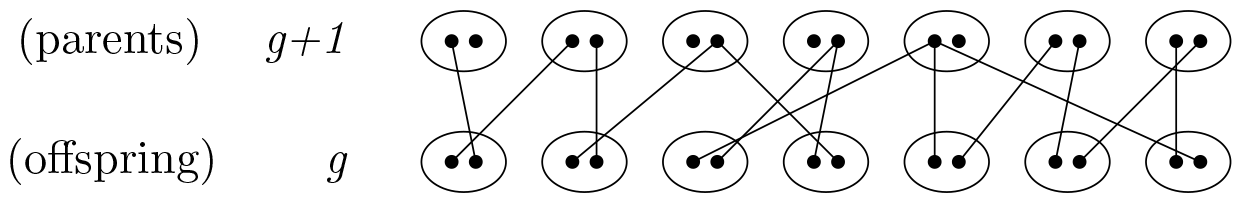
2. With probability *α*_*N*_, a pair of adults is chosen uniformly without replacement to have a very large number of offspring, [*ψN*] where *ψ* ∈ [0, 1] is a fixed fraction of the population. The other *N* − [*ψN*] offspring are produced as above according to the Wright-Fisher model. In this case we call *g* a “*generation with a big family*”. An example of this special reproduction dynamics is depicted below for *N* = 7 and *ψ* = 0.72 in which the highly reproductive pair (I_1_, I_2_) = (4, 5) in generation *g* + 1 has [*ψN*] = [0.72 *·* 7] = 5 offspring in generation *g*.

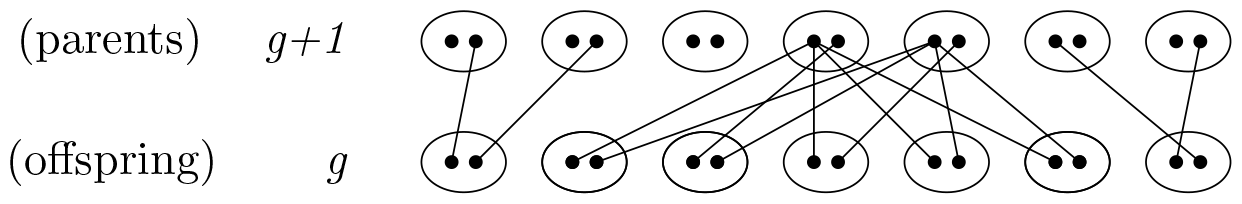

These two possibilities happen independently for all generations *g* ∈ *ℤ*_≥0_. The classical Wright-Fisher model corresponds to the case when *α*_*N*_ = 0. In this case every *g* ∈ *ℤ*_≥0_ is a Wright-Fisher generation. Note, we allow selfing with probability 1/*N* for all offspring produced by Wright-Fisher reproduction but we assume that the [*ψN*] offspring of big families have two distinct parents.

The *parent assignment* between (parental) generation *g* + 1 and (offspring) generation *g* is the collection of edges connecting the offspring with their parents. The diagram in (1) below shows the parent assignment corresponding to the example above in which *g* is a Wright-Fisher generation.

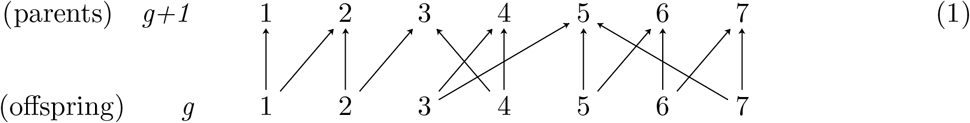

On the other hand, the diagram in (2) below shows the parent assignment corresponding to the example above in which *g* is a generation with a big family.

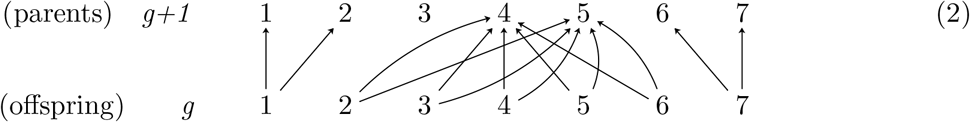

### Pedigree

The collection of all the parent assignments among all pairs of consecutive generations is called the *pedigree* and it is denoted as 𝒜^(*N*)^. The pedigree models the set of all family relationships among the members of the population for all generations. The pedigree is shared among all loci. It is the structure through which genetic lineages are transmitted. Patterns of ancestry, or gene genealogies are outcomes of Mendelian inheritance in this single shared pedigree.

### Frequency of big families

Recall that *α*_*N*_ denotes the probability of a big family to appear in a generation. We set

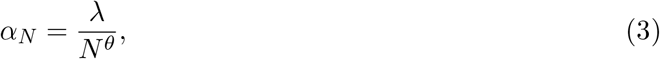

where *θ* ∈ (0, 1] and *λ* ∈ *ℝ*_≥0_ is a fixed parameter which determines the relative frequency of big families on the timescale of *N*^*θ*^ generations.

### Timescale

Suppose two individuals are sampled uniformly *without* replacement among the *N* individuals of the current generation *g* = 0 and we sample one gene copy from each. Let *τ*^(*N*,2)^ be the pairwise coalescence time, that is, the number of generations in the past until the two sampled gene copies coalesce. How long is the pairwise coalescence time *τ*^(*N*,2)^? This will depend on *N* and also on *θ* owing to our assumption (3). In considering the limiting ancestral process for the sample, we re-scale time so that it is measured in units of *N*^*θ*^ generations. We study the distribution of the re-scaled pairwise coalescence time, *τ*^(*N*,2)^/*N*^*θ*^, with different results depending on whether *θ* ∈ (0, 1) or *θ* = 1. In the latter case, our timescale is *N* generations, which we note is 1/2 the usual coalescent timescale for diploids. In the former case, where we may infer from (3) that big families will dominate the ancestral process, the timescale is accordingly much shorter than the usual coalescent timescale. Coalescence times in both cases also depend on a combined parameter *ψ*^2^/4 which is the limiting probability of coalescence when a big family occurs.

### Limiting process by averaging over the pedigree

For reference and to illustrate our choice of timescale, we begin with a Kingman coalescent approximation for the pairwise coalescence time in the classical Wright-Fisher model, here the special case *θ* = 1 and *λ* = 0 or *α*_*N*_ = 0. Averaging over the process of reproduction in a single generation gives a coalescence probability of 1/(2*N*). With *θ* = 1, we measure time in units of *N* generations. To parallel the derivation of our main result, we consider the probability that the coalescence time *τ*^(*N*,2)^ is more than [*tN*] generations. The limiting ancestral process is obtained as

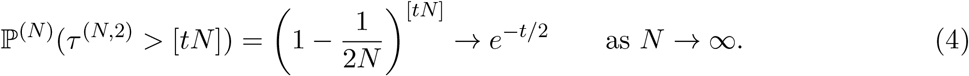

In words, the re-scaled coalescence time 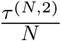 converges in distribution to an exponential random variable with rate parameter 1/2.

Before stating our main result, we first prove Lemma 1 below, generalizing (4) to our population model, in the sense that for *θ* = 1 with *ψ* = 0 or *λ* = 0 in Lemma 1 we recover (4).

#### Lemma 1.

*Let λ* ∈ *ℝ*_≥0_, *θ* ∈ (0, 1], *and set* 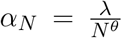. *The re-scaled coalescence time*^*τ*^ 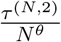 *converges in distribution to an exponential random variable with rate parameter*

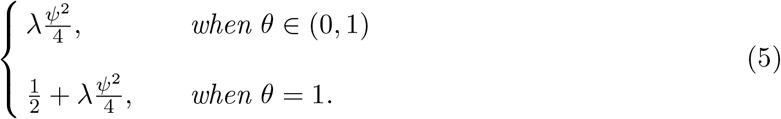

We note that in Birkner et al. (2013b), the full ancestral recombination graph for samples of arbitrary size and genomes consisting of arbitrary numbers of linked loci is described for a population model nearly identical to ours here. The ancestral recombination graph (Hudson, 1983a; Griffiths and Marjoram, 1997), like the Kingman coalescent itself, averages over the pedigree. Lemma 1 describes the marginal ancestral process for a sample of size two at a single locus.

#### Proof of Lemma 1

The lineage dynamics of our model can be analyzed using a Markov chain. In any generation *g* in the past, the ancestral lineages of a pair of gene copies must be in one of the three states {*ξ*_0_, *ξ*_1_, *ξ*_2_}, where

- *ξ*_0_ = 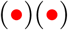 represents two ancestral lineages in two distinct individuals,
- *ξ*_1_ = 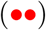 represents two ancestral lineages on different chromosomes in the same individual,
- *ξ*_2_ = 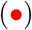 represents that the ancestral lineages have coalesced.

The diploid ancestral process for a pair of gene copies can thus be represented as a Markov chain 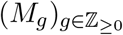 with state space {*ξ*_0_, *ξ*_1_, *ξ*_2_}, where *M*_*g*_ is the state of the two lineages *g* generations in the past. Its one-step transition matrix *Π*_*N*_ is given by

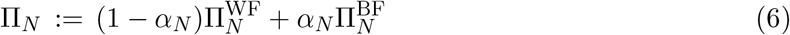

where

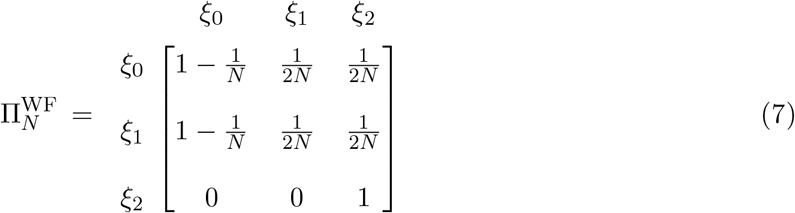

and

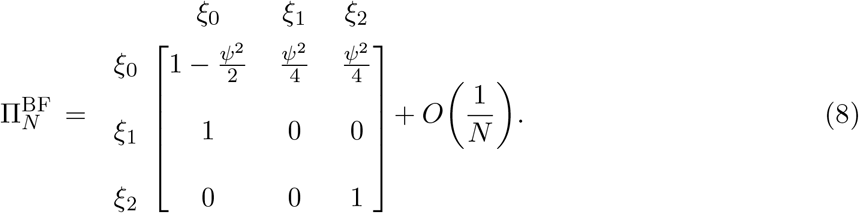

The matrix 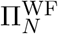 in (7) is the transition matrix for a Wright-Fisher generation, whereas 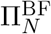 in (8) is for a generation with a big family. The entries of 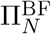 in (8) are derived by conditioning on the parent assignment(s) for the individual(s) containing the ancestral lineages, with respect to the highly reproductive pair. For instance, for ancestral lineages currently in two distinct individuals, the coalescence probability is 1/4 if both individuals are members of the big family and 1/(2*N*) otherwise. Thus we have

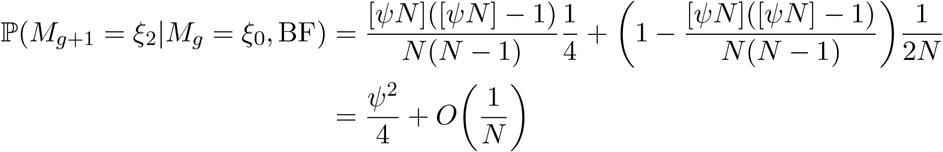

for the transition *ξ*_0_ → *ξ*_2_ in 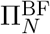, and where we have also specified that this contribution to the overall probability in (6) is conditional on the occurrence of a big family.

The rest of the proof is a straightforward application of Möhle (1998a, Lemma 1). This is a separation-of-timescales result. To see how it works, using (3) we can rewrite (6) as

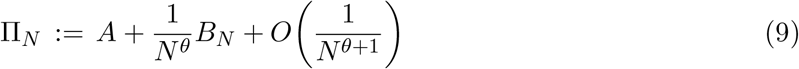

where

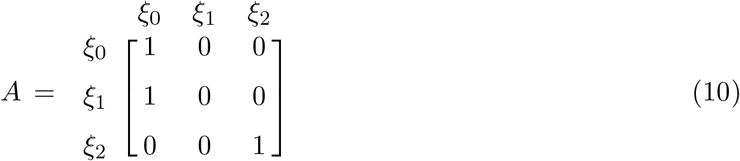

and

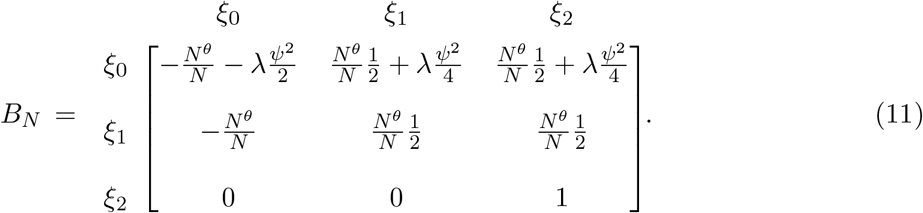

The matrix *A* contains the fastest parts of the process. The matrix *B*_*N*_ contains the next-fastest parts of the process, specifically those occurring on the timescale of *N*^*θ*^ generations.

Möhle’s result depends on the existence of equilibrium stochastic matrix *P* := lim_*k*→*∞*_ *A*^*k*^ which in this case is equal to *A*. Möhle’s result also requires the existence of the limiting infinitesimal generator *G* := lim_*N*→*∞*_ *PB*_*N*_ *P*. Note that in our application *B* := lim_*N*→*∞*_ *B*_*N*_ itself converges. From (11) it is clear that the limiting result will differ depending on whether *θ* ∈ (0, 1) or *θ* = 1. If *θ* ∈ (0, 1), the contribution of Wright-Fisher generations to the coalescence rate shrinks to zero in the limit. If *θ* = 1, the contribution of Wright-Fisher generations, which is 1/2 on our timescale, remains comparable to the contribution of generations with big families in the limit.

Applying Möhle (1998a, Lemma 1) to compute our probability of interest,

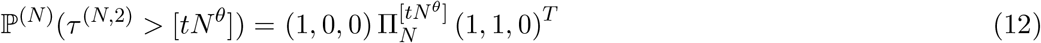

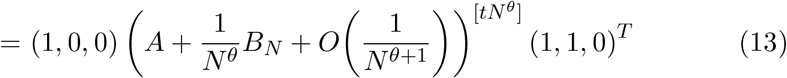

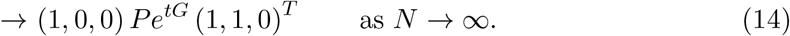

The initial vector (1, 0, 0) enforces our assumed starting state, *ξ*_0_. The end vector (1, 1, 0)^*T*^ enforces the requirement that the lineages remain distinct at generation [*tN*^*θ*^], i.e. that the Markov chain 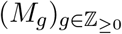 has not reached state *ξ*_2_. Möhle’s result *Pe*^*tG*^ is in the middle. Recall that *P*, which here is equal to *A*, instantaneously adjusts the sample so that the effective starting state is *ξ*_0_ even if the sample state is *ξ*_1_. The lineages then enter the continuous-time process with rate matrix *G*. Overall we have

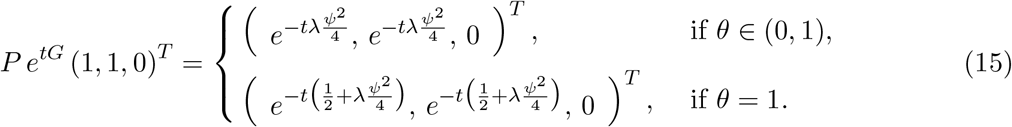

The right hand side of (14) is equal to (5), and the proof of Lemma 1 is complete. □

#### Remark 1

(Robustness against perturbation of initial condition). The form of *P* shows that the limiting result in Lemma 1 holds regardless of whether the sample begins in state *ξ*_0_, as we have assumed, or in state *ξ*_1_. So, other sampling schemes could be considered. In fact Lemma 1 still holds if the initial distribution lies in the set ℐ := {(c, 1 − c, 0) ∈ [0, 1]^3^ : c ∈ [0, 1]}. This can be seen clearly in (15).

### Limiting process by conditioning on the pedigree

Our main result is about the conditional distribution. We let

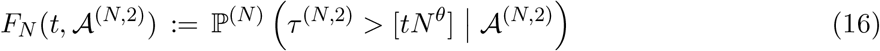

be the conditional probability of the event {*τ*^(*N*,2)^ > [*tN*^*θ*^]} given the (random) pedigree and the sampled pair of individuals. Mathematically, 𝒜^(*N*,2)^ is the sigma-field (all information) generated by the outcome of the random reproduction of the population and the knowledge which pair of individuals was sampled.

#### Theorem 1.

*Let λ* ∈ *ℝ*_≥0_, *θ* ∈ (0, 1], *and set* 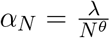. *For all t* ∈ (0, *∞*), *we have the following convergence in distribution as N* → *∞*

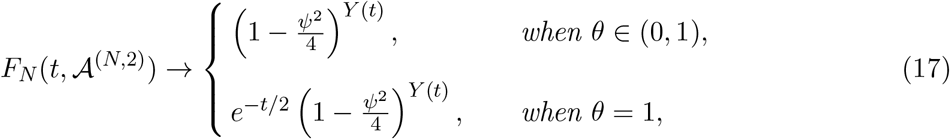

*where Y* (*t*) *is Poisson process with rate λ*. *In fact, the convergence in* (17) *holds jointly for all t* > 0, *see the discussion in Remark 4 in the Appendix section A*.*4 for details*.

Theorem 1 offers a description of the conditional distribution of the coalescence time *τ*^(*N*,2)^ for a sample of two genes in a population of size *N* given the pedigree. It says that the law of 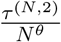, under the conditional probability ℙ(*·* | *𝒜*^(*N*,2)^), converges weakly as *N* → *∞* to the law of a random variable (call it *T*) under a probability measure ℙ_*Y*_ that depends on the Poisson process *Y* with rate *λ*. Furthermore, the survival function ℙ_*Y*_ (*T* > *t*) is equal to the right hand side of (17). In what follows, we will refer to F_*N*_ (*t*, 𝒜^(*N*,2)^) defined in (16) as the discrete survival function.

Theorem 1 has an intuitive interpretation. Taking the case *θ* = 1, the *e*^−*t/*2^ represents the probability that the two lineages have not coalesced by time *t* due to ordinary Wright-Fisher/Kingman coalescence. Against this smooth backdrop there are *Y* (*t*) points, representing essentially instantaneous events in which a big family occurs and the lineages have a large probability, *ψ*^2^/4, of coalescing. Thus there is an additional factor in the survival function representing the probability that the pair does not coalesce in any of these extreme events. The case *θ* ∈ (0, 1) is analogous except the timescale is so short that there is no chance of an ordinary Wright-Fisher/Kingman coalescent event.

Note that when *λ* = 0, there are no large reproduction events and *Y* (*t*) ≡ 0. Then for *θ* ∈ (0, 1), the right hand side of (17) is 1, i.e. there is no coalescence with probability 1. For *θ* = 1, the right hand side of (17) is *e*^−*t/*2^ which is expected from the cumulative distribution function (CDF) of the Kingman coalescent for a sample of size 2, with our timescale. The degenerate case *λ* > 0 but *ψ* = 0 effectively gives these same results for any *Y* (*t*).

### Proof of Theorem 1

Recall that each *g* ∈ *ℤ*_≥0_ is a Wright-Fisher generation (resp. a generation with a big family) with probability 1 − *α*_*N*_ (resp. *α*_*N*_), independently for all *g* ∈ℤ_≥0_. The number of generations with big families in {0, 1, …, *G*−1}, denoted by *H*_*N*_ (*G*), therefore has the binomial distribution Bin(*G, α*_*N*_).

We begin by addressing the technical point that we cannot actually know just by looking at the pedigree whether *g* is a generation with a big family, the way we have defined these as occurring only in special generations. Even in the classical Wright-Fisher model, every individual has the capacity to produce a large number of offspring. But reproductive outcomes as extreme as our big families are exceedingly rare under ordinary Wright-Fisher reproduction when *N* is large.

To illustrate, consider the event that, spanning generations *g* + 1 and *g*, there exists a pair of parents with at least [*ψN*] offspring. In our population model, this is guaranteed to occur in generations with big families. Note that the two parents of a big family have an additional ∼Poisson(2(1 − *ψ*)) offspring because the other *N* − [*ψN*] offspring are produced according to the Wright-Fisher model. The event that a pair of parents with at least [*ψN*] offspring can also occur randomly in Wright-Fisher generations, but only with small probability

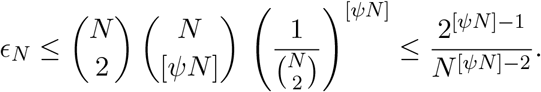

Let *Q*_*N*_ (*G*) be the number of generations *g* ∈ {0, 1, …, *G*−1} in which such an event occurs between *g* + 1 and *g*. Then *Q*_*N*_ (*G*) is extremely close to the binomial variable *H*_*N*_ (*G*) ∼ Bin(*G, α*_*N*_) because

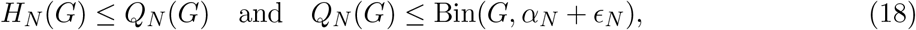

where the first inequality holds almost surely and the second is a stochastic dominance. Since 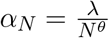, for each *t* ∈ (0, *∞*) and *θ* ∈ (0, 1] we have convergence in distribution

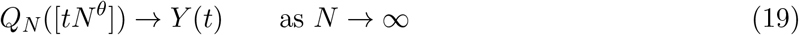

which is identical to the limiting result for *H*_*N*_ ([*tN*^*θ*^]). In other words, ϵ_*N*_ is so small for any sizeable *N*, that we are safe in assuming that such extreme events in the pedigree reliably signify generations with big families as defined under our model.

Indeed, from the discussion above we have

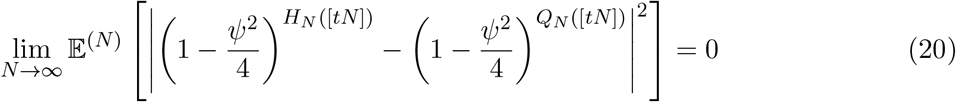

so that we can (and will) in the following computations replace *H*_*N*_ ([*tN*]) by *Q*_*N*_ ([*tN*]) without changing any limit as *N* → *∞*.

**Proof of** (17) **when** *θ* = 1 In this case it suffices to show that

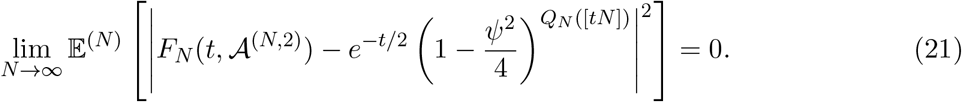

Expanding the square in (21) gives

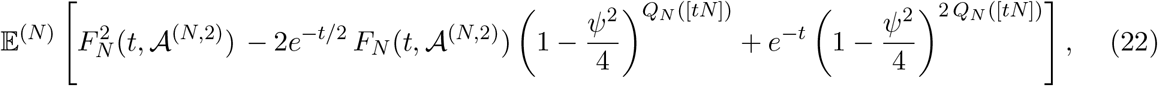

which requires the computation of three expectations. The first is the expectation of the square of the discrete survival function,

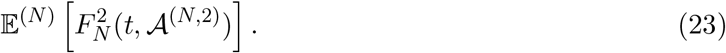

The second is the expectation of the discrete survival function times the probability that a single pair of lineages does not coalesce in any of the generations with big families in the pedigree up to time *t*,

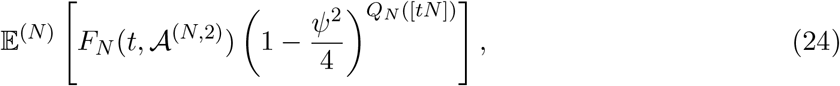

The third is the expectation of the square of the same, latter probability that a single pair of lineages does not coalesce in any of the generations with big families in the pedigree up to time *t*,

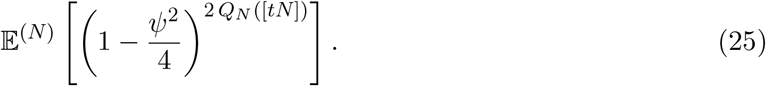

**First term in** (22) The expectation in (23) can be computed by considering two samples of size 2 whose lineage dynamics are conditionally independent given 𝒜^(*N*,2)^. Genetically, this corresponds to the ancestral processes of two unlinked loci given the pedigree and the two sampled individuals, and where one gene copy has been sampled at each locus from each of the individuals. Let *τ* and *τ*^*′*^ be the coalescence times of these two pairs of sampled gene copies. Due to the conditional independence of these coalescence times, for all *g* ∈ *ℤ*_+_ we have

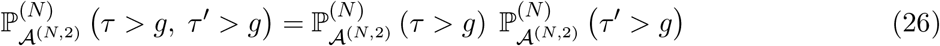

in which 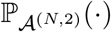 is short-hand for (ℙ *·*|*𝒜*^(*N*,2)^) in (16). Setting *g* = [*tN*] and taking expectations on both sides of (26) gives

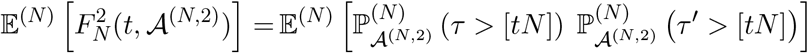

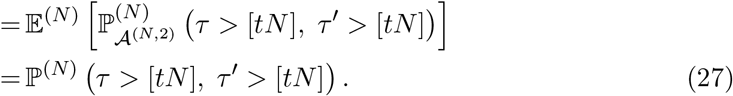

In order to compute the limit as *N* → *∞* in (27), we introduce the ancestral process of two conditionally independent samples given the pedigree.

### Joint diploid ancestral process

The stochastic dynamics of the two conditionally independent, given the pedigree, pairs of lineages are described by the *joint diploid ancestral process* 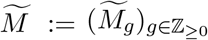. This is a Markov chain with state space 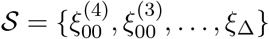 described below, where 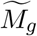 is the state of the two pairs of lineages in a common pedigree *g* generations backwards in time. Denote by 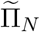 its transition matrix, the derivation of its entries is available at A.1.1 and its entries are available at A.1.2 for a generation with a big family and at A.1.3 for a Wright-Fisher generation.

Similarly to the proof of Lemma 1, denote by 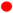 an ancestral lineage of a gene copy in the first pair and by 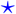 the same for the second pair. Parentheses are used to denote individuals. More precisely, consider the following 10 states:

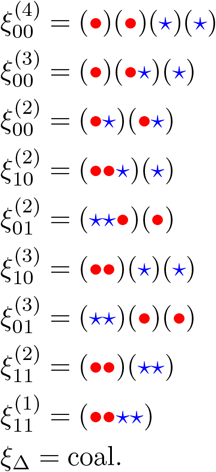

The superscript indicates the total number of individuals in which the 4 ancestral lineages reside. The two subscripts tell us the states of the two pairs respectively: 0 means a pair of lineages in state *ξ*_0_ and 1 means a pair of lineages in state *ξ*_1_, with these as defined in the proof of Lemma 1. For example, 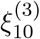 involves 3 individuals in which the first pair of lineages are in the the same individual and the second pair of lineages is in different individuals. Finally, the state *ξ*_Δ_ is an absorbing state which represents the event that at least one of the two pairs has coalesced. The order of the states is arbitrary, based first on the subscripts then on the superscripts.

By definition, the two pairs of gene copies are drawn from the same pair of individuals at the present generation *g* = 0, where for each pair one gene copy is picked from each of the individuals. Hence the initial state 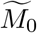 must be 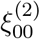. In other words, the distribution 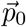 of 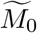 is given by

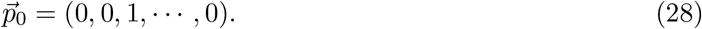

It follows from Lemma 2 in Appendix Section A.1.4 that

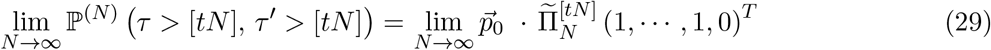

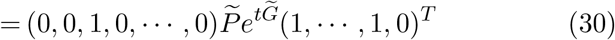

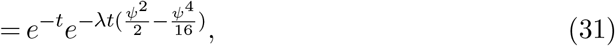

where (29) follows from the definition of 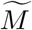 and 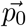, (30) from (Möhle, 1998a, Lemma 1) as explained in Section A.1.4 and (31) by Lemma 2. Note that the vector (1, *· · ·*, 1, 0)^*T*^ in (29)-(30) amounts to the Markov chain 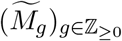 not reaching state *ξ*_Δ_, i.e. that neither pair has coalesced.

#### Remark 2

(Robustness of joint process to initial condition). Our assumed initial state 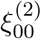 is the usual way multi-locus data are sampled in population genetics. But Lemma 2 and Theorem 1 both hold for any initial state 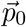 whose last coordinate is zero. This is because the sample will undergo an instantaneous adjustment by 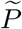 given in (A20), so that the effective starting state is always 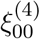. Whatever idiosyncrasies 𝒜^(*N*,2)^ may possess, especially in the recent past, sensu Chang (1999), meaning the most recent log_2_(*N*) generations, these matter less and less as *N* grows. In the limit, the lineages of any sample immediately disperse to different individuals without undergoing any coalescent events. Similarly, the factors of 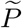 in 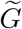 guarantee that the lineages will remain in state 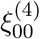 throughout the ancestral process, except for instants in which they have a chance to coalesce. This robustness against initial condition is analogous to (14).

**Second term in** (22) We now show that (24) converges to 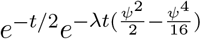 as *N* → *∞*. Through the use of the law of total expectation, (24) is equal to

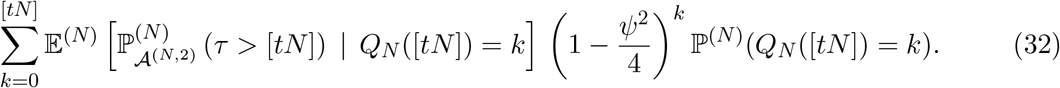

By the fact that *Q*_*N*_ ([*tN*]) is known given the pedigree and an application of the tower property, the conditional expectation in (32) is equal to

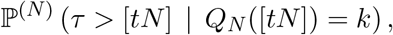

which is approximately equal to

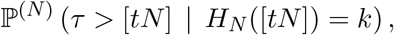

by (18). That is to say

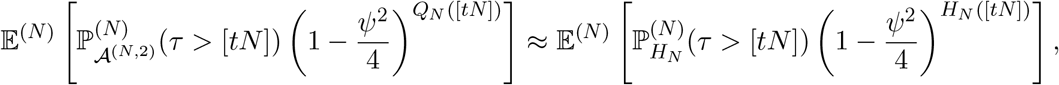

in the sense that (20) holds. By Lemma 3 in the Appendix, for *g* = [*tN*], it follows that for each *N* ≥ 2 and *t* ∈ (0, *∞*),

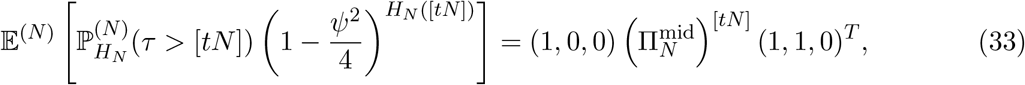

where 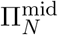 is defined as

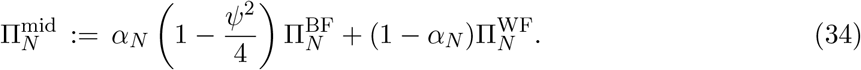

It now follows by Lemma 4 that the right hand side of (33) converges to 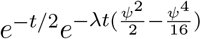.

**Third term in** (22) Finally, (25) is computed by first noticing that the number of big families up to generation *T, Q*_*N*_ (*T*), is (almost) binomially distributed according to Bin([*T* ], *α*_*N*_) for all *T* ≥ 1, as observed by (18). Using the probability generating function of *Q*_*N*_ (*T*) we get that the third term in (22) is equal to 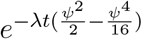.

### Putting everything together

As *N* → *∞*, (22) is equal to 0 since (25) multiplied by *e*^−*t*^ and (23) add up to 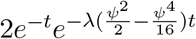 which cancel out with (24) multiplied by −2*e*^−*t*^. This gives (21) which concludes the proof of Theorem 1 in the case of *θ* = 1.

**Convergence** (17) **when** *θ*∈ (0, 1) The proof is similar to the case of *θ* = 1. In all of the above, substitute [*tN*] by [*tN*^*θ*^], and show instead of (22) that

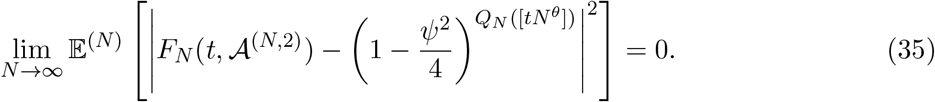

Expanding (35) gives the same three terms as in (23)-(25). In this faster timescale, as *N* → *∞*, (23) is now equal to

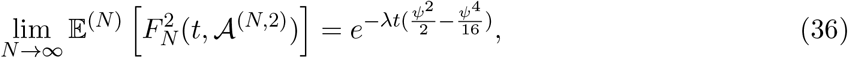

as available in Lemma 2. The limiting behavior of (24) is the same as before, that is

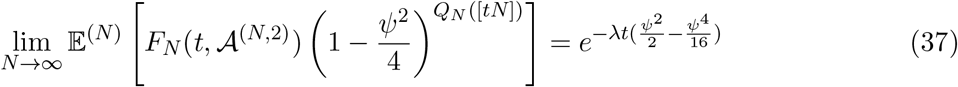

and

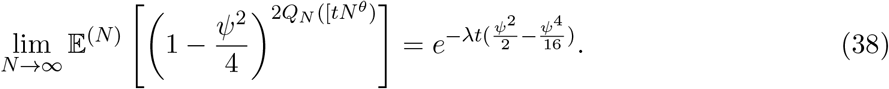

Multiplying (37) by −2 and summing it up with (36) and (37) concludes the proof in the case of *θ* ∈ (0, 1). □

The proof of Theorem 1 is complete.

#### Remark 3

(Only big families matter). Let 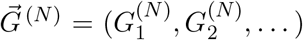, where 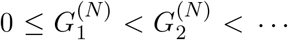, be the generations with big families that have (randomly) occurred. Then 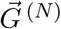 is known if we know the pedigree. Similar to (16), we let

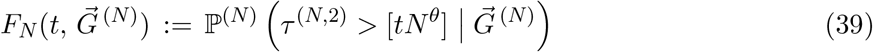

be the conditional probability of the event {*τ*^(*N*,2)^ > [*tN*^*θ*^]} given the (random) generations 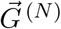. Hence, here we condition on less information than on the left hand side of (17). We can show that Theorem 1 still holds (i.e. the weak convergence in (17) still holds) if we replace F_*N*_ (*t*, 𝒜^(*N*,2)^) by 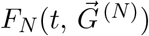. For a proof sketch see Appendix A.2.1.

### Coalescence times, gene genealogies and correlations

Here we briefly recap then provide three illustrations of our results. Our main result is Theorem 1 which describes two limiting distributions of coalescence times conditional on the pedigree. As the number of unlinked loci examined in the sampled individuals increases, the empirical distribution of their coalescence times should converge to Theorem 1. In this case, conditional on the pedigree, the probability of coalescence in a generation depends on whether that particular generation includes a big family. For background and comparison, Lemma 1 presents the corresponding two limiting distributions obtained by the usual method of averaging over pedigrees, i.e. over all possible outcomes of reproduction in a single generation, including the possibility of a big family. In this case, the probability of coalescence is the same in every generation.

Time is re-scaled in all of these limiting ancestral processes. It is measured in units of *N*^*θ*^ generations for some *θ* ∈ (0, 1]. When *θ* ∈ (0, 1), the timescale for big families to occur is much shorter than the usual Wright-Fisher coalescent timescale of *N* generations. When *θ* = 1, the timescales for big families and for ordinary Wright-Fisher coalescence are the same. Big families occur at rate *λ* in re-scaled time, and their offspring comprise a fraction *ψ* ∈ [0, 1] of the population in that generation. Underpinning our results is the fact that as *N* → *∞* ancestral genetic lineages spend the overwhelming majority of their time in separate individuals, i.e. in state *ξ*_0_ for a pair of lineages at the same locus (cf. Lemma 1) or state 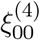 for two pairs of lineages at two unlinked loci (cf. Theorem 1 and Remark 2). Thus when a big family occurs, each lineage independently: (i) is among the offspring of the highly reproductive pair with probability *ψ* and (ii) if so, is equally likely to descend from each of the four copies of the corresponding locus in the two parents. A pair of lineages at the same locus coalesces in the big family with probability *ψ*^2^/4. Pairs of lineages at different, unlinked loci do this independently.

Our first illustration compares our limiting results to the cumulative distribution function (CDF, i.e. one minus the survival function) of pairwise coalescence times in the discrete model. Figure 1a displays CDFs for five simulated pedigrees for *N* = 500, assuming that the probability of a big family is equal to the expected pairwise coalescence probability, 1/(2*N*) = 0.001, and the offspring make up the entire population in that generation. This corresponds to the limiting process in Theorem 1 with *θ* = 1, *λ* = 1/2 and *ψ* = 1. This makes the coalescence probability (*ψ*^2^/4) equal to 1/4 in each generation with a big family. We computed coalescence probabilities on each pedigree in each generation starting from a pair of randomly sampled individuals using the method in Wakeley et al. (2012). The corresponding “expected” CDF of the pedigree-averaged process from Lemma 1, i.e. of an exponential random variable with rate parameter 5/8, is shown for comparison.

**Figure 1:**
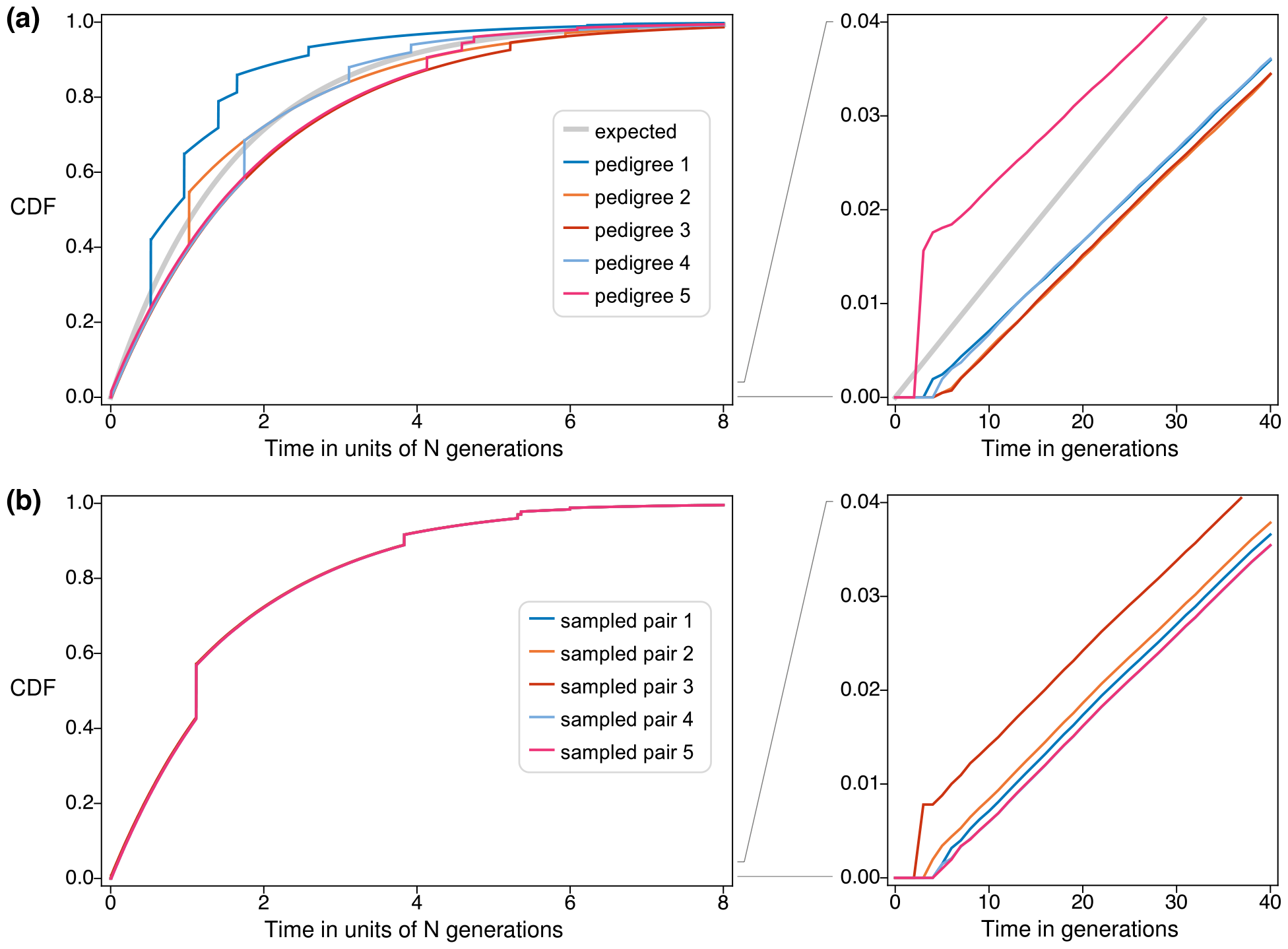
Cumulative distribution functions (CDFs) of pairwise coalescence times for *θ* = 1 and *λ* = 1/2. (a), left panel: CDFs for five simulated pedigrees for populations of size *N* = 500 together with the corresponding expected CDF from Lemma 1. (a), right panel: The same five CDFs and the corresponding expectation from Lemma 1, only plotted over the most recent 40 generations. (b): corresponding results for a single pedigree for a population of size *N* = 500 but five different pairs of individuals, each sampled independently without replacement from the population.

The left panel of Figure 1a illustrates that the ancestral process conditional on the pedigree is quite close to limiting result in Theorem 1, even when *N* = 500. The CDFs make discrete jumps whenever big families occur. In this case with *ψ* = 1 the magnitude of a jump is always 1/4 of the remaining distance to 1. Between jumps the CDFs show a steady increase in the cumulative coalescence probability, in line with the limiting prediction with its rate of 1/2. In contrast, the pedigree-averaged process in Lemma 1 predicts a faster rate of increase of the CDF and no jumps.

The right panel of Figure 1a details the short-time behavior of the ancestral process conditional on the pedigree, displaying these same CDFs only over the most recent 40 generations. The scale on the vertical axis is such that the diagonal corresponds approximately to the prediction of the background Wright-Fisher model (not shown) and a line with slope 1.25 corresponds approximately to prediction of Lemma 1 which is shown. After a small number of generations, which from Chang (1999) should be of order log_2_(*N*), the CDFs for the five pedigrees start to show the predicted

Wright-Fisher slope of one. However, they start at different places depending on the particular ancestries of the sampled individuals, specifically whether there are very recent shared ancestors as in pedigree 5 or more likely there are no very recent shared ancestors as in pedigrees 1 through 4; cf. also Wakeley et al. (2012). These differences are barely visible on the timescale of the left panel of Figure 1a, and it is implicit in Theorem 1 that they become negligible as *N* → *∞*.

As stated in (26), the predictions for each of the five pedigrees in Figure 1a apply equally and independently to every locus in the sampled individuals. These five, like five instances of Theorem 1, are again predictions for the empirical distributions of coalescence times among unlinked loci. Different instances of 𝒜^(*N*,2)^ will have different times of big families (Figure 1a, left panel) and different patterns of recent common ancestry of the samples (Figure 1a, right panel). For comparison, Figure 1b shows the same two graphs for five independently sampled pairs of individuals on a single pedigree. Again, each sample has its own pattern of recent common ancestry, producing visible differences on the scale of the right panel. But now all five samples access the same shared set of big families, resulting in the five closely overlapping CDFs in the left panel of Figure 1b.

Next we illustrate the effects that big families have on the gene genealogies of larger samples, in particular the sharing of identical coalescence times at unlinked loci. Rather than simulating pedigrees for finite populations, we use the limiting model directly so that big families are the only possible cause of shared coalescence times. We set *ψ* = 1 as before, and for simplicity assume that big families drive the ancestral process, i.e. *θ* ∈ (0, 1). We set *λ* = 1 without loss of generality, as *λ* is arbitrary except when *θ* = 1.

Based on Theorem 1, we model gene genealogies by generating a series of exponential waiting times between big families and, since *θ* ∈ (0, 1), disallowing coalescence between them. When the *n* ancestral lineages of the sample reach the first big family, their distribution among the four parental gene copies will be multinomial with parameters *n* and (1/4, 1/4, 1/4, 1/4). Anywhere from one to four simultaneous multiple mergers will occur. The number of ancestral lineages which emerge is also at most four. If more than one lineage emerges, the same process is repeated until a single lineage remains which is the most recent common ancestor of the entire sample. The only aspects of the pedigree which persist in the limit are the big families (cf. Remark 3). Thus, independent runs of this multinomial coalescent process using the same series of exponential waiting times correspond to gene genealogies of unlinked loci conditional on the pedigree.

Figure 2a displays the gene genealogies of seven unlinked loci for a sample of size 16, assuming in this way that all loci share the same pedigree. The trees are oriented with the present-day samples at the bottom. Solid lines trace (unlabeled) ancestral lineages up into the past. Thin dotted lines show the times of the big families. All seven gene genealogies have multiple-mergers at the most recent big family in the past, and five have common ancestor events at the second one. In the more distant past when there are small numbers of ancestral lineages, there is less sharing of coalescence times among gene genealogies. This is expected; for example, the final two lineages only coalesce with probability 1/4 each time they encounter a big family.

**Figure 2:**
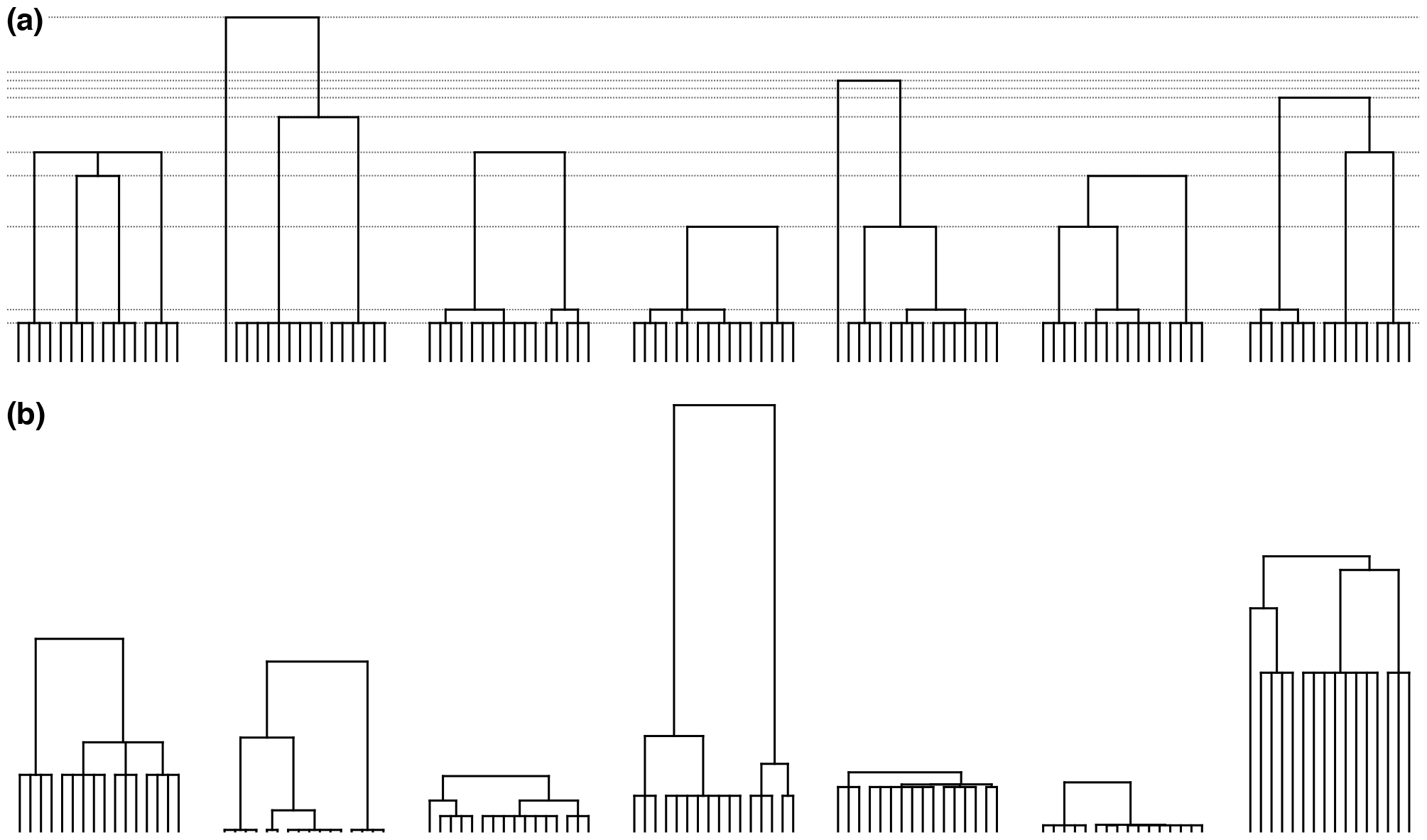
Simulated gene genealogies for seven independently assorting loci when all seven share the same pedigree (a) versus when each locus has its own independently generated pedigree (b). The sample size is *n* = 16 for every locus. Gene genealogies were generated as described in the text for the limiting model with *θ* ∈ (0, 1) and *λ* = *ψ* = 1. Thin dotted lines in the top row show the particular series of times of big families in that population.

Figure 2b shows seven gene genealogies, again for samples of size 16, but now assuming that each locus has its own pedigree. These are equivalent to seven gene genealogies sampled from seven independent populations, each with its own series of exponential waiting times between big families as in Figure 2a (not displayed in Figure 2b). These gene genealogies differ from the ones in the top row, most obviously in the different timings of their first common ancestor events. Clearly, the distribution of gene genealogies produced in this way will not be close to the distribution of gene genealogies of unlinked loci in the same genome which perforce come from the same population.

Finally, we illustrate how averaging over pedigrees as in Lemma 1 results in positive correlations of coalescence times between unlinked loci. Explicitly modeling pedigrees as in Theorem 1 predicts these to be zero as might be expected for independently assorting loci. Based on the property of ancestral lineages spending the overwhelming majority of their time in separate individuals, cf. Remarks 1 and 2, we consider the two-locus analogue of Lemma 1 with reduced state space

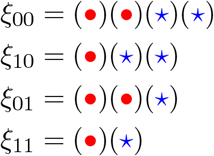

where now the first and second subscripts are indicators of whether locus 1 or locus 2 has coalesced. By extension from Lemma 1, the limiting ancestral process for two unlinked loci has transition rate matrix

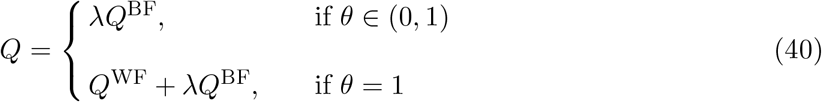

where

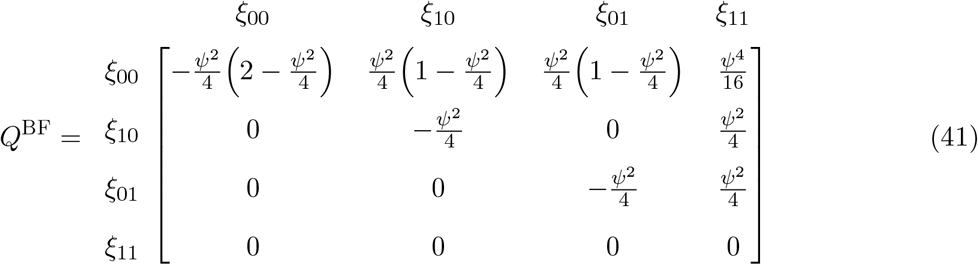

and

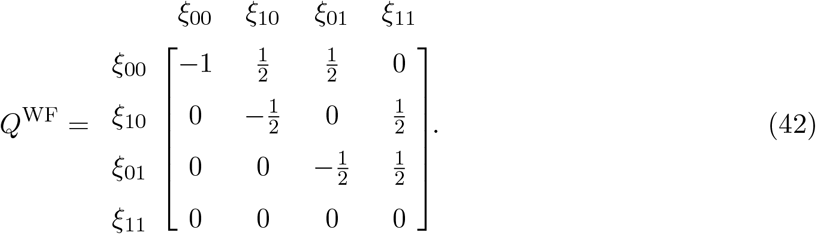

Focusing on the case *θ* = 1, the rate matrix *Q* is the sum of a Wright-Fisher (or Kingman coalescent) component and a big-family component. We have factored the tuning parameter *λ* out of the latter to emphasize that, conditional on the occurrence of a big family, samples at the two loci coalesce or do not coalesce independently of each other.

Let *T*_1_ and *T*_2_ be the coalescence times at the two loci. These correspond to the limiting random variables *τ*/*N*^*θ*^ and *τ*^*′*^/*N*^*θ*^ in Lemma 2. Here individually they are the times to state *ξ*_11_ starting from states *ξ*_01_ and *ξ*_10_, respectively. From the rate matrix *Q* in (40) or from Lemma 1 directly, *T*_1_ and *T*_2_ are identically distributed. In particular,

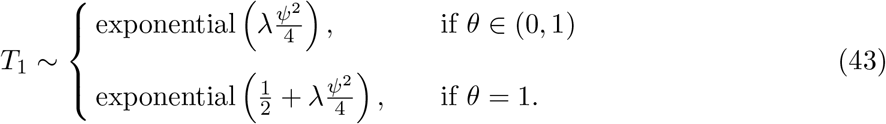

However, *T*_1_ and *T*_2_ are not necessarily independent. Lemma 2 accounts for this non-independence in the proof of Theorem 1, and we note that (40) also gives (A22). Here, we use first-step analysis to compute the correlation coefficient, Corr[*T*_1_, *T*_2_]. Let W be the waiting time to the first event in the ancestry of the two loci starting from state *ξ*_00_, and 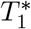 and 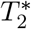 be the additional times to coalescence at each locus following the first event. In this formulation,

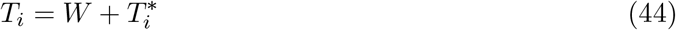

for *i* ∈ {1, 2}. From (40), we have

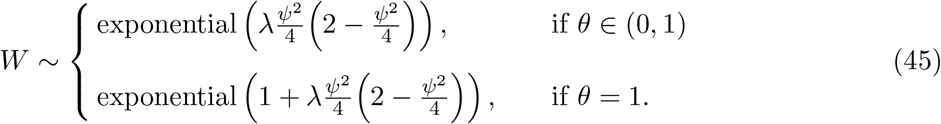

and we point out that, since *W* is exponentially distributed,

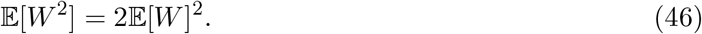

Conditioning on the first step from state *ξ*_00_ and simplifying,

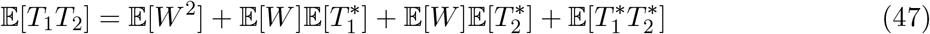

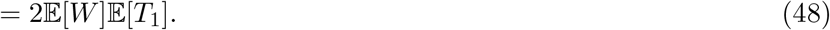

Going from (47) to (48) uses (46), (44), 𝔼 [*T*_1_] = 𝔼 [*T*_2_], and the fact that either 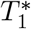 or 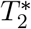 or both are equal to zero following the first event. Then for the correlation coefficient, we have simply

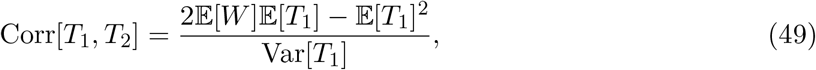

which, using (43) and (45), becomes

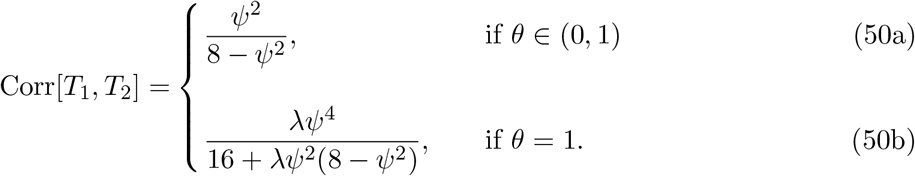

Even though the loci assort independently, their ancestries in the pedigree-averaging model jointly depend on the random process that generates big families in the population. As a result, their coalescence times are positively correlated.

The correlation coefficient (50b), obtained here under the assumption that the loci are unlinked, corresponds to Equation 31 in Birkner et al. (2013b, p. 266), obtained there by modeling recombination explicitly and then taking the limit as the re-scaled recombination parameter tends to infinity. The timescales in these two works differ by a factor of two. Our (50b) becomes identical to Equation 31 in Birkner et al. (2013b) by putting *λ* = c/2.

For a given value of *ψ*, the correlation coefficient is smaller when *θ* = 1 than when *θ* ∈ (0, 1). When coalescence can be due to either big families or ordinary Wright-Fisher reproduction (*θ* = 1), the correlation tends to zero as *λ* tends to zero. As *λ* grows, (50b) grows until it approaches (50a). Thus, the occurrence of big families may be said to be the source of positive correlations in coalescence times at unlinked loci. In a similar vein, Corr[*T*_1_, *T*_2_] tends to zero as the fraction of the population replaced by each big family, *ψ*, tends to zero. This is true even if *θ* ∈ (0, 1), i.e. when there is no Wright-Fisher/Kingman component in the limit process. At the other extreme, as *ψ* → 1, Corr[*T*_1_, *T*_2_] → 1/7 which is considerably less than one. Even when all coalescence happens in big families and the offspring of each big family replace the entire population, there are still two diploid parents and the loci will generally have different coalescence times.

The following alternate derivation of (50a) shows how these positive correlations arise. In short it is because *T*_1_ and *T*_2_ have a shared dependence on the times between big families in the pedigree. Implicitly, Lemma 1 averages over these times whereas Theorem 1 retains them.

When *θ* ∈ (0, 1), coalescence can only happen when a big family occurs. Let *K*_1_ and *K*_2_ be the numbers of such events it takes for locus 1 and locus 2 to coalesce, respectively. These do not depend on the times between big families when *θ* ∈ (0, 1). Further, *K*_1_ and *K*_2_ are independent because the loci are unlinked. They are geometric random variables with parameter *ψ*^2^/4. Let *X*_*i*_, *i* ∈ *ℤ*_≥0_, be the time from the (*i* − 1)th to the *i*th big family backward in time, with *X*_0_ ≡ 0. In the context of Theorem 1, these times are independent and identically distributed exponential random variables with rate parameter *λ*. Under this formulation,

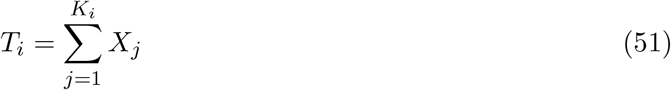

for *i* ∈ {1, 2}. There are two sources of variation in *T*_*i*_: variation in *K*_*i*_ and variation in the lengths of the intervals, *X*_*j*_, j ∈ {1, …, *K*_*i*_}. Starting with (51), it is straightforward to confirm that the distribution of *T*_*i*_ is exponential with rate parameter *λψ*^2^ /4 as in (43) or Lemma 1.

From (51) and the fact that *X*_*i*_ and *X*_*j*_ are independent for *i* ≠ *j*, it is also clear that intervals in a common to *T*_1_ and *T*_2_ are a key source of their covariation. For given values of *K*_1_ and *K*_2_, they are the only source. The first interval is always shared, as are all subsequent intervals until one or the other locus coalesces. Let *K*_12_ be the number of these shared intervals and

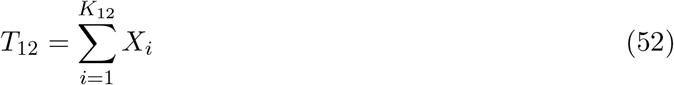

be the corresponding total length of time. By definition *K*_12_ = min(*K*_1_, *K*_2_). The more ancient *K*_1_ − *K*_2_ or *K*_2_ − *K*_1_ intervals are only ancestral to one of the loci.

Applying the conditional covariance formula, or law of total covariance, we have

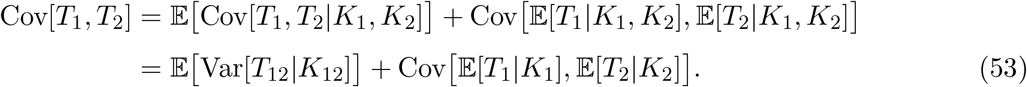

The outer expectation and covariance are with respect to the joint distribution of *K*_1_ and *K*_2_. Note that *K*_12_ is a marginal property of this distribution. The inner variance (or covariance) and expectations are with respect to the joint distributions of the *X*_*i*_ which are the only parts of *T*_1_ and *T*_2_ that vary conditional on *K*_1_ and *K*_2_.

At this point, in (53), we have not applied the fundamental property that *K*_1_ and *K*_2_ are independent since the loci are unlinked, nor have we assumed any particular distribution(s) for the *X*_*i*_. We have only used the definitions of *T*_1_ and *T*_2_ as sums of random variables and the assumption that *X*_*i*_ and *X*_*j*_ are independent for *i* ≠ *j*. So we may consider that the interval times are fixed numbers: *X*_*i*_ ≡ *x*_*i*_, *i* ∈ *ℤ*_≥0_. They could be the outcomes of the exponential random times implicit in Theorem 1. Fixing the *X*_*i*_ means fixing the only aspects of the pedigree that persist in the limiting model. Conditioning on the pedigree, *T*_1_ and *T*_2_ are independent even in the limiting model; cf. (26). The point we wish to emphasize here is that fixing the *X*_*i*_ removes one particular source of covariation of *T*_1_ and *T*_2_. It makes Var[*T*_12_|*K*_12_] = 0.

Continuing from (53) and assuming that *X*_*i*_, *i* ∈ *ℤ*_≥0_, are independent and identically distributed

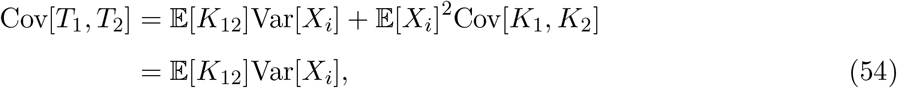

the latter following from the independence of *K*_1_ and *K*_2_. Again, *X*_*i*_ ∼ exponential(*λ*), and from the definition of *K*_12_ as the number of big-family events it takes for one locus or the other to coalesce,

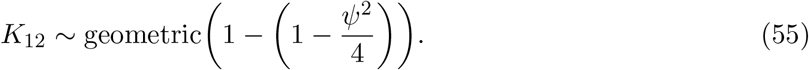

Putting the required quantities in (54) and simplifying gives

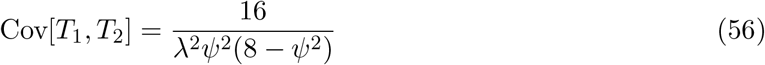

which is exactly the covariance needed to produce the correlation coefficient (50a). In sum, the model of Lemma 1 predicts a positive correlation of coalescence times at unlinked loci because it averages over the distributions of the intervals *X*_*i*_. Starting instead with the model of Theorem 1 shows that the particular quantity controlling these positive correlations is Var[*X*_*i*_].

## Discussion

The use of random models to describe past events raises many questions in population genetics. Everything in the past has already occurred, including all instances and timings of reproduction and genetic transmission. For empirical work this may be a truism. But population genetics has always been concerned with evolutionary processes. How do mutation, recombination, selection, random genetic drift, non-random mating, limited dispersal, etc., conspire to produce observable patterns of variation? By emphasizing the fixed nature of the past, we highlight the subjectivity of theoretical work, specifically when the goal is to interpret data from natural populations. Ultimately, the choices one makes about modeling the past may be application-dependent.

Motivated by applications to multi-locus data, we singled out the pedigree as a key feature of the past and obtained a result (Theorem 1) concerning the application of neutral coalescent models in sexually reproducing species. We have the following sampling structure in mind. Processes of survival and reproduction result in a pedigree. Genetic transmission, including mutation and recombination across the entire genome, occurs within the pedigree. A number of individuals are sampled from the population and some or all of their genomes are sequenced. We modeled the single-locus coalescent process conditional on the pedigree. Our results specify the distribution of coalescence times given the pedigree and the sampled individuals. This distribution can be interpreted either as a prior for a single locus or as a prediction about the distribution of coalescence times among unlinked loci. We contrasted our results conditional on the pedigree with results obtained by averaging over pedigrees, noting that the latter is the tradition of theoretical population genetics. We did not model mutation or recombination, but our fundamental conclusion—that some population processes cause the quenched and averaged processes to be very different—should be as important for genetic variation as it is for coalescence times.

We can compare our framework with that of Ralph (2019). The two have a lot in common. Ralph (2019) takes the pedigree and the outcomes of genetic transmission, including recombination across the entire genome, to be fixed. The latter is referred to as the ancestral recombination graph (ARG), which we note differs slightly from the corresponding objects in Hudson (1983a) and Griffiths and Marjoram (1997) because it is embedded in the fixed pedigree. Without specifying a generative model for the pedigree, Ralph (2019) focuses on the ARG as the fixed but unknown object of interest in empirical population genetics. A sample is taken and some stretch of the genome is sequenced. Its ancestry is a collection of gene genealogies, a subset of the ARG. Implicitly, it is the outcome of the random process of genetic transmission within the fixed pedigree, but this too is not modeled.

The only randomness is in how the collection of gene genealogies of the sample is revealed by mutation. Ralph (2019) assumes the infinite-sites mutation process and uses this to show that predictions about summary statistics of DNA sequence variation, such as the average number of pairwise nucleotide differences or the F-statistics of Patterson et al. (2012), can be expressed in terms of the fixed branch lengths in the sampled subset of the ARG. This is the empirical version of what Slatkin (1991), Griffiths and Tavaré (1998), Nielsen (2000), McVean (2002) and Peter (2016) had done in the context of the standard neutral coalescent, where instead the moments of summary statistics can be expressed in terms of corresponding moments of branch lengths. Ralph et al. (2020) describe a hybrid approach, with the ARG conceived as in Ralph (2019) and with times of events in the ARG for data from humans (The 1000 Genomes Project Consortium, 2015) estimated with the aid of the standard neutral coalescent (Speidel et al., 2019).

Whereas we model the production of the pedigree and the process of coalescence within it but do not model mutation, Ralph (2019) models only mutation on the fixed ARG. Consider a species in which recurrent selective sweeps across the genome have structured the ARG. An empirical estimate of the ARG would find regions of the genome with reduced variation due to reduced times to common ancestry. In order to relate these observations to an evolutionary process within the empirical framework of Ralph (2019), for example to describe them in terms of recurrent selective sweeps as in Durrett and Schweinsberg (2005), additional modeling would be needed. In contrast, in a theoretical approach such as ours here, recurrent sweeps would be included in the model at the outset, and this in turn would facilitate the interpretation of patterns in the data. Under our model, it is important to keep in mind that the ARG is in fact a fixed object and that the process of coalescence within the pedigree models the sampling of a locus in the ARG.

Today detailed estimates of the ARG for large samples of human genomes are available (Wohns et al., 2022; Zhang et al., 2023). These have been obtained, like other recent estimates (Kelleher et al., 2019; Speidel et al., 2019; Albers and McVean, 2020), using the standard neutral coalescent as a prior for gene genealogies and times to common ancestry. Our results and those of Tyukin (2015) help to justify using such a prior despite the fact that the pedigree is fixed, so long as the processes which laid down the pedigree are not too different from the Wright-Fisher or Cannings models with relatively low variation of offspring numbers. The empirically oriented interpretations in these works, for example in Wohns et al. (2022), connect features of the ARG with major events in human history, such as the out-of-Africa event which has been studied genetically since the first mtDNA discoveries (Cann et al., 1987; Vigilant et al., 1991) and the novel finding of an accumulation of ancestry in Papua New Guinea more than 100-thousand years ago. This is intraspecific phylogeography (Avise et al., 1987; Avise, 1989, 2000) at genome scale.

Our model for generating the pedigree includes the possibility of special generations in which a big family is guaranteed to occur. We obtained different coalescent processes as *N* → *∞*, depending on the relative rate of these big families in the limit and whether the ancestral process is conditional on the pedigree (Theorem 1) or not (Lemma 1). This essentially negative result, that the averaged process cannot be used in place of the conditional process, includes the positive finding that the Kingman coalescent can be used between big families in the case that both occur on the same timescale (Theorem 1 with *θ* = 1). The numbers and timings of big families are all that is left of the pedigree in the limit (cf. Remark 3). Needing to keep track of just these is much less daunting than the prospect of including entire pedigrees in all of our population-genetic models. There may be other circumstances in which aspects of the pedigree are important, but so far the only other instance identified is when sub-populations are connected by limited migration (Wilton et al., 2017).

Limiting coalescent processes for our model generally involve simultaneous multiple-mergers. Yet the familiar extensions of the Kingman coalescent to include multiple-mergers have been derived by averaging over the pedigree, not by conditioning on it. They begin with single-generation marginal probabilities of coalescence, whereas in truth the individuals in the sample either have or do not have common ancestors in any preceding generation and this is what determines probabilities of coalescence. Without big families, our results and those of Tyukin (2015) provide belated justification for the early uses of the Kingman coalescent process as a prior model for the gene genealogy of a single locus (Lundstrom et al., 1992; Griffiths and Tavaré, 1994; Kuhner et al., 1995). Our work also clarifies what is involved in using pedigree-averaged ancestral processes as single-locus priors in cases where big families can occur.

If the only data available were from a single locus without recombination, one could model the gene genealogy using the pedigree-averaged ancestral process. The logic would be that a single locus has one unknown random pedigree and one unknown random gene genealogy within that pedigree, and that the gene genealogies from multiple-mergers coalescent models are marginal predictions over both of these unknowns. For example, with *n* = 2, a single draw of a coalescence time from the appropriate exponential distribution in Lemma 1 accounts for both sources of variation. Implicit in this accounting is that repeated samples would each have their own pedigree and conditional gene genealogy. This two-fold sampling structure is precisely what Theorem 1 describes. It is straightforward to show that repeated sampling under Theorem 1 (each time drawing a new pedigree) gives the same exponential distributions as in Lemma 1. Yet even for a single locus, it may be preferable to record the additional information about big families as in Theorem 1.

Applying this type of repeated sampling (i.e. including re-sampling the pedigree) to multiple loci is another matter. Population-genetic models should not allow the pedigree to vary among loci. Theorem 1 is a simple initial example of the kind of coalescent modeling required for multi-locus data generally but especially when multiple-mergers processes are implicated. In cases where big families may occur with some frequency, it is crucial to retain the information about the pedigree which matters for the gene genealogies at all loci. All multiple-mergers coalescent models so far described, which implicitly average over the pedigree, are inadequate in this sense. The broader implication of Lemma 1 and Theorem 1 is that there exists a collection of quenched limits conditional on the pedigree which await description and are the appropriate models for multi-locus data.

The diploid exchangeable population models in Birkner et al. (2018) are a natural starting point for the description of general quenched-pedigree Ξ-coalescent models. Alternatively, parameterized models could be considered, controlling for example rates of monogamy and the distribution of offspring numbers as in the program SLiM 3 (Haller and Messer, 2019) or the Pólya urn scheme of Gasbarra et al. (2005). The latter was used for the prior in the Bayesian inference methods of Gasbarra et al. (2007a,b) and Ko and Nielsen (2019) for estimating the recent few generations of the pedigree from sequence data. Selfing in the production of big families, which we assumed does not occur, could also be considered. Non-exchangeable models for generating pedigrees are possible, for example with recurrent selective sweeps (Durrett and Schweinsberg, 2005) or cultural transmission of reproductive success (Guez et al., 2023). It could also be of interest to describe these coalescent models directly in terms of the properties of pedigrees as directed graphs, and here we note the study of Blath et al. (2014) as a start in this direction.

The model underlying Lemma 1 and Theorem 1 is very simple. Only one type of big family is allowed, these are distributed in time according to a Poisson process, and we only considered a sample of size two. We hypothesize that the basic principles of Theorem 1 will be robust to all of these. For example, other ways of generating big families should be possible, such that *Y* (*t*) ∼ Poisson(*λt*) would be replaced by some other distribution. Extensions to larger sample sizes and to variation in the numbers and types of big families seem straightforward in principle, though they will require a lot more bookkeeping. Such generalization of big families will need to include sufficient details that Mendel’s laws can be applied. For example, if four parents have [*ψN*] offspring, it will matter whether they form two monogamous pairs or comprise one big family with four parents, and in either case just how many offspring each pair has. In a more general model, such details will need to be specified for each big-family event.

The implications of our results for inference can be sketched as follows. Consider a general situation in which there may be special events like our big families in a well mixed population which has possibly changed in size over time. Assume that data are available for *L* unlinked loci and there is no intra-locus recombination. Let 𝒟, 𝒢, 𝒜, Θ_m_ and Θ_c_ represent the data, the collection of gene genealogies at the loci, the pedigree, the parameters of a mutation model and the parameters of a coalescent model, specifically a trajectory of relative population sizes over time in a Kingman coalescent with variable size. Let 𝒟_*i*_ and 𝒢_*i*_ be the data and the gene genealogy at the *i*th locus. Consider the likelihood, which is key to any sort of statistical inference. Traditional coalescent-based inference disregards 𝒜 and computes the likelihood ℙ(𝒟; Θ_m_, Θ_c_) in which we use “;” to indicate that Θ_m_ and Θ_c_ are treated as fixed parameters. The traditional computation proceeds by conditioning on 𝒢, treating this as a random variable, but does so under the erroneous assumption that ℙ(𝒢; Θ_c_) is equal to the product of ℙ(𝒢_*i*_; Θ_c_) across loci.

Instead, because a shared pedigree has been fixed by past events, a better approach would be to use 𝒜 as the parameter in place of Θ_c_ and to compute the likelihood

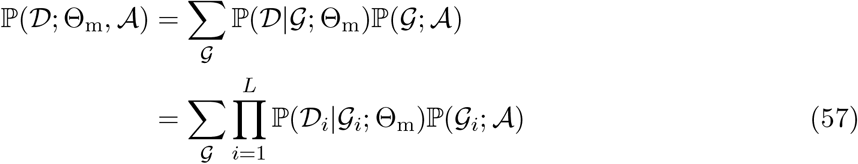

where now in (57) the independence assumption of gene genealogies (given the pedigree) is correct. Theorem 1 is a simple example of what we expect will be possible under a variety of population models. Intuitively, *𝒜* can be replaced by the pair {*𝒴, 𝒜 \ 𝒴*}, where *𝒴* is a list of special events and *𝒜 \ 𝒴* is the remainder of the pedigree. In the limiting ancestral process, *𝒴* may need to be preserved while *𝒜\ 𝒴* can be replaced by a coalescent model with parameters Θ_c_. For our model, *𝒴* would be the times and sizes (*ψ*) of big families, and the coalescent model would be the Kingman coalescent. Thus our results suggest the simplification

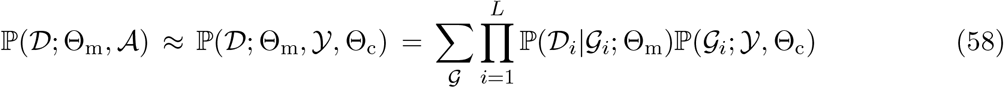

where the approximation is for large *N*. In the present work, (58) has the probabilistic interpretation in Theorem 1, where *𝒜* and the limiting object *𝒴* are random outcomes of a population process. Then ℙ(*𝒴*) could also serve as the prior for Bayesian inference of *𝒴*, using (58) but with “|” not “;” for conditioning on *𝒜* and *𝒴*. In any case, the pair {*𝒴*, Θ_c_} is a much more manageable variable than *𝒜*. For many species, it will not be necessary to record special events in the limiting model. Without *𝒴*, (58) reduces to traditional coalescent-based inference.

The issues we raise here about pedigrees parallel those in recent work on population bottlenecks. Like the trajectories of population sizes through time in coalescent hidden Markov models, bottlenecks have traditionally been considered fixed events of the past. But models of recurrent bottlenecks have recently been considered. A bottleneck is the event that a population ordinarily of size *N*_0_ has size *N*_*B*_ < *N*_0_ for a period of time. In Birkner et al. (2009, Section 6) it was shown that a Ξ-coalescent describes the limiting gene-genealogical process for a model with recurrent severe bottlenecks, specifically with the bottleneck duration going to zero and *N*_*B*_/*N*_0_ → 0 as both *N*_*B*_ and *N*_0_ go to infinity. González Casanova et al. (2022) used a similar framework but allowed that *N*_*B*_ could be finite. They described a new class of Ξ-coalescents they called the symmetric coalescent. A model like ours with *ψ* = 1 and random selfing between the parents of the big family would give one of these, being identical to a short drastic bottleneck (González Casanova et al., 2022, Definition 3) with *N*_*B*_ = 4 and our *θ* and *λ* corresponding to their *α* and k^(*N*)^. But as noted in Birkner et al. (2009), Ξ-coalescent models are only obtained for recurrent bottlenecks by averaging over the exponential process which generates them. When the times and severities of bottlenecks are fixed, the result will depend on these and will not be a time-homogeneous Markov process. Against this backdrop of similarities, a small but notable difference is that the bottleneck models in Birkner et al. (2009) and González Casanova et al. (2022) are haploid rather than diploid.

The proof of Theorem 1 uses the idea of two independent copies of the coalescent process on the same pedigree. Genetically, these are the gene-genealogies of two independently assorting loci conditional on their shared pedigree. Our results suggest a reinterpretation of population-genetic models which predict non-zero correlations or positive covariances of coalescence times at unlinked loci. Eldon and Wakeley (2008) found that the correlation could be positive in a model of recombination and the haploid (or gametic) equivalent of big families. Birkner et al. (2013b) extended this finding to a diploid model of recombination with big families similar to the ones we studied here. It appears that such correlations result from averaging over the pedigree. In the simple model we considered, they arise from averaging over the times of big families, cf. (53) and (54). On any fixed pedigree the correlation of coalescence times at unlinked loci must be zero.

The comparison with recurrent bottlenecks is apt here as well. Schaper et al. (2012) constructed a recurrent-bottleneck model for recombination and coalescence at two loci with recombination. They note that what is relevant for data is the covariance of coalescence times conditional on the series bottleneck events in the ancestry of the population, not the unconditional covariance which averages over these. They showed that the conditional covariance goes to zero as the recombination parameter goes to infinity. They also showed, in the Ξ-coalescent limit of Birkner et al. (2009) that the covariance could be positive even as the recombination parameter goes to infinity. We note an analogous finding for yet another model in Wakeley and Lessard (2003), in which non-zero correlations of coalescence times at unlinked resulted from taking the number of subpopulations to infinity in an island migration model, even though for any finite number of subpopulations the correlation goes to zero as the recombination parameter tends to infinity.

In sum, for a century it has been common practice in population genetics to compute probabilities of past events by averaging over an assumed process of reproduction. What we have shown is that when big families occur with some frequency, or more generally when the descendants of a small number of individuals take over a sizable fraction of the population in a short period of time, this averaging is not justified and can produce spurious results. Instead, such extreme outcomes of reproduction should be viewed as fixed, and probabilities of coalescence and other events conditioned upon them. In light of this, existing multiple-mergers coalescent models must be reassessed and most likely replaced with conditional or quenched models. A comparison with how population size has been treated as fixed is of some interest because it too is an outcome of reproduction. In both cases, it is when population-genetic models are applied to explain variation among loci that the importance of conditioning on past events is most readily apparent.

## Data availability

Mathematica notebooks containing some calculations, detailed in the Appendix, are available at: https://github.com/diamantidisdimitris/Bursts-of-coalescence.

## Acknowledgements

We thank Bjarki Eldon for helpful discussions, and two anonymous reviewers for insightful comments. This work was support in part by National Science Foundation grants DMS-1855417 and DMS-2152103, and Office of Naval Research grant N00014-20-1-2411 to Wai-Tong (Louis) Fan. Dimitrios Diamantidis is supported by NSF DMS-2152103.

## A Appendix

### A.1 Joint diploid ancestral process

Recall the joint ancestral process 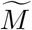 describes the lineage dynamics of two conditionally independent samples of size 2. In section A.1.1 we establish its Markov property. This property should hold for any population model in which the outcomes of reproduction in different generations are independent. In section A.1.2 we provide its transition matrix for a generation with a big family and in section A.1.3 we study two extreme cases, *ψ* = 0 and *ψ* = 1. In the former case we recover the transition matrix of the joint diploid ancestral process for a Wright-Fisher generation and in the latter we consider the case when the highly reproductive couple replaces the entire population.

#### A.1.1 The joint diploid ancestral process with 10 states is Markovian

Recall the state space of the process 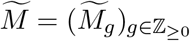 is 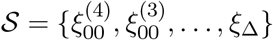 which has 10 states including the state *ξ*_Δ_ where one, the other or both pairs of lineages have coalesced.

Fix a pair of states (*ξ*, η) ∈ *𝒮*^2^. It holds that

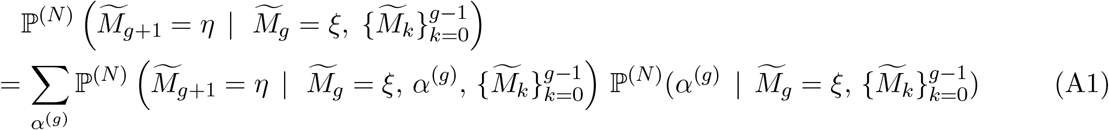

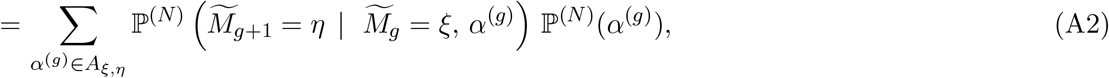

where the last sum 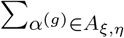 is over all possible parent assignments *α*^(*g*)^ that belong to the pattern *A*_*ξ,η*_ described in Tables A1-A3. These are exactly the parent assignments for which transition to η is *possible* in the sense that 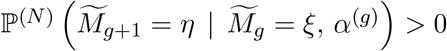 if and only if 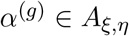.

In the last equality above, we noted that a parent assignment *α*^(*h*)^ between generations *g* and *g* + 1 is independent of 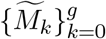 and so 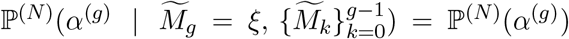. Also, 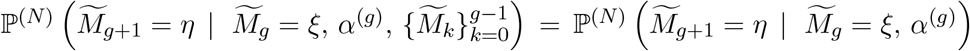 because it is determined entirely by Mendelian randomness and not by the population model (whether it is Wright-Fisher or something else). This conditional probability is computed in Tables A4-A10, and we only need to compute this for those *α*^(*g*)^ in patterns whose probability is of order 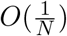 or larger in view of Möhle (1998a, Lemma 1).

In particular, we have shown that the number 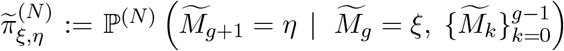 does not depend on the “history” 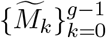. Hence the process 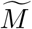 is Markovian.

#### A.1.2 Transition matrix for a generation with a big family

For any *ξ* ∈ *𝒮*, let 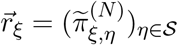 be the *ξ*th row of 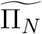. By (A2), each row of 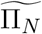 is computed as

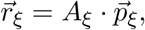

where *A*_*ξ*_ is given by the appropriate matrix in Tables A4-A10 corresponding to *ξ*, and 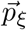 contains the probabilities of the patterns from Tables A1-A3 used in all the transitions from *ξ* to η ∈ *𝒮*. In a generation with a big family, 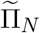 is given as

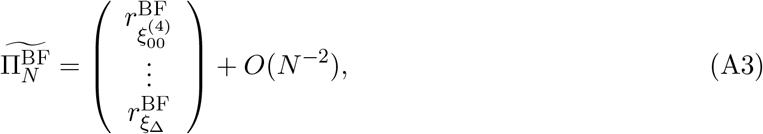

where 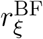 in (A3) is the row of 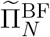 that corresponds to initial state *ξ* up to an error of order O(*N*^− 2^), given in (A4)-(A8). For example, 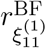 is derived by multiplying the matrix 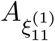 given in Table A4 by the vector column 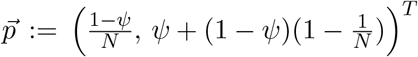 on the right. The calculations for each row are available at https://github.com/diamantidisdimitris/Bursts-of-coalescence in the **PiBF_row_derivation_print** file and can be reproduced in the corresponding Mathematica notebook **PiBF_row_derivation_print_notebook**, where Mathematica version 13.1.0 (Wolfram Research, Inc., 2022) was used. Its rows are given as

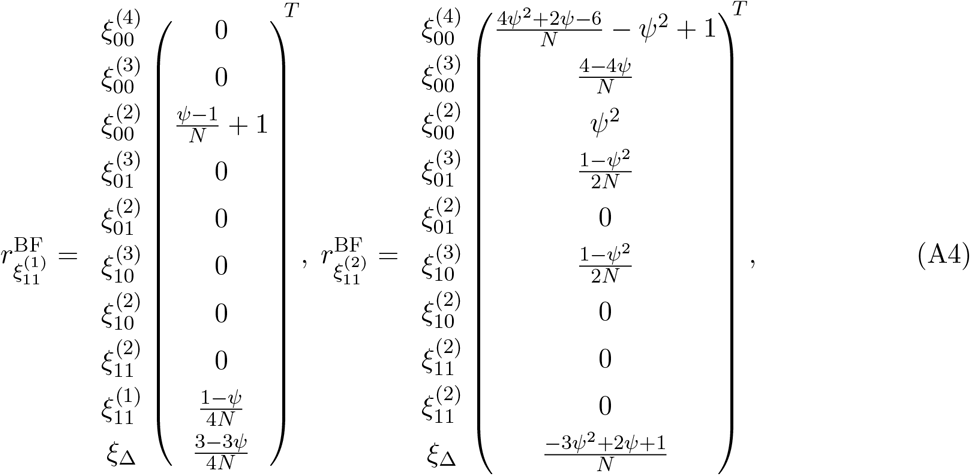

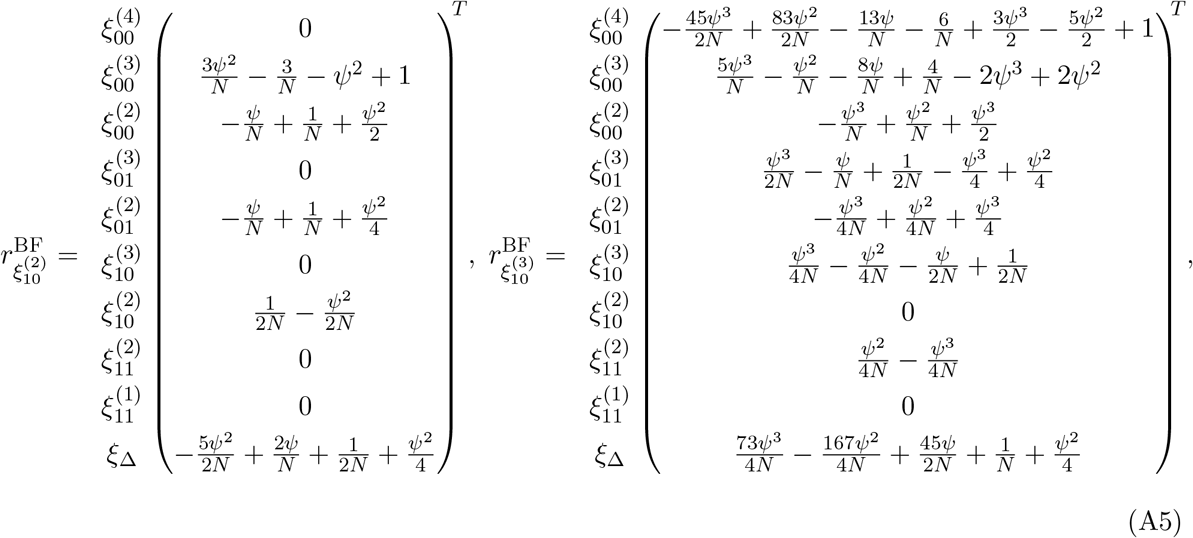

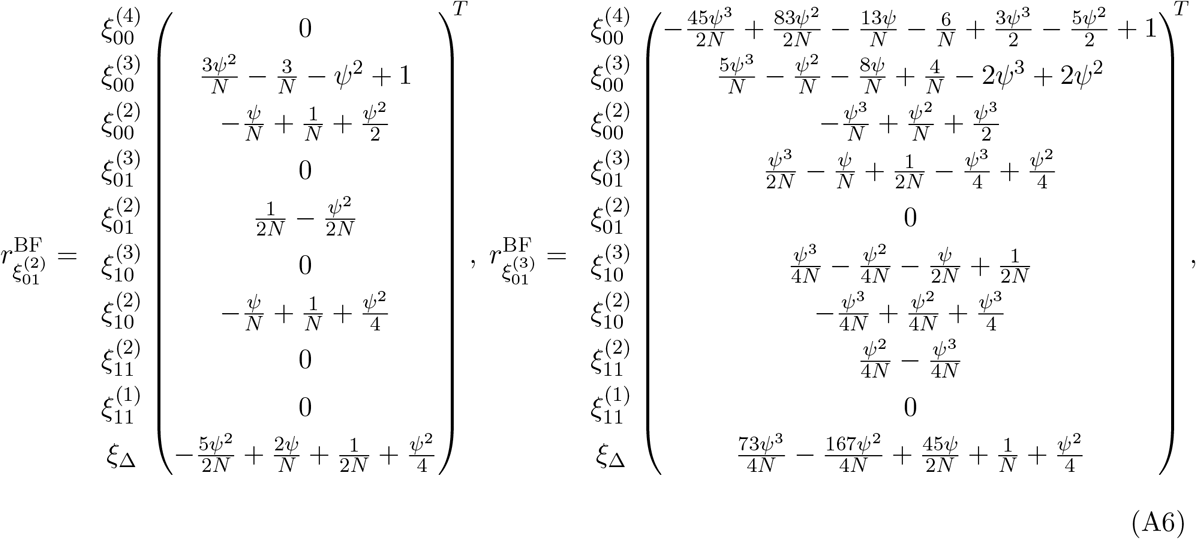

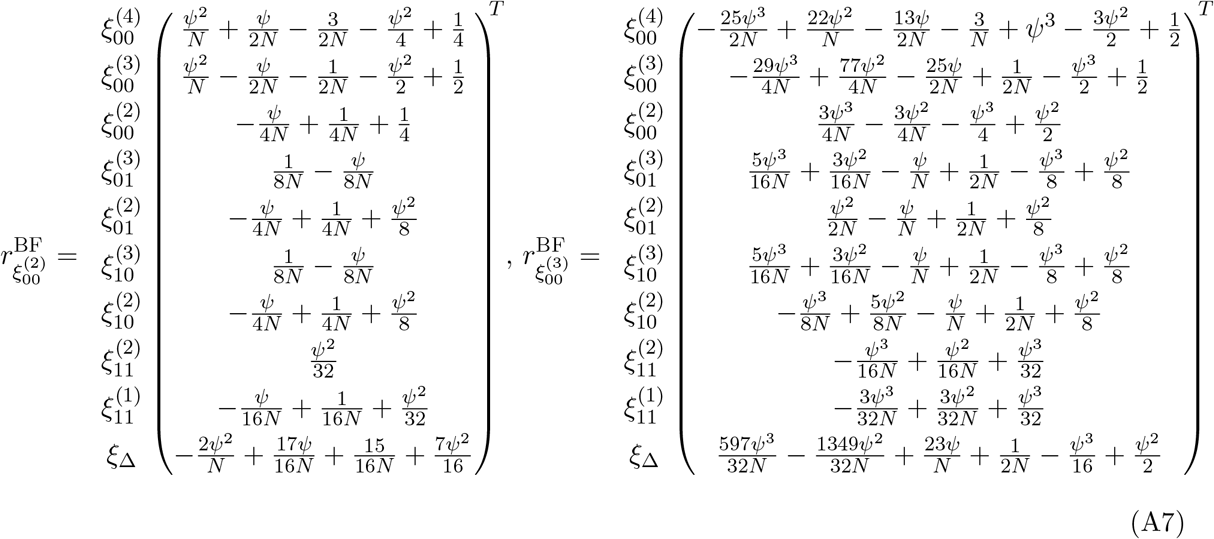

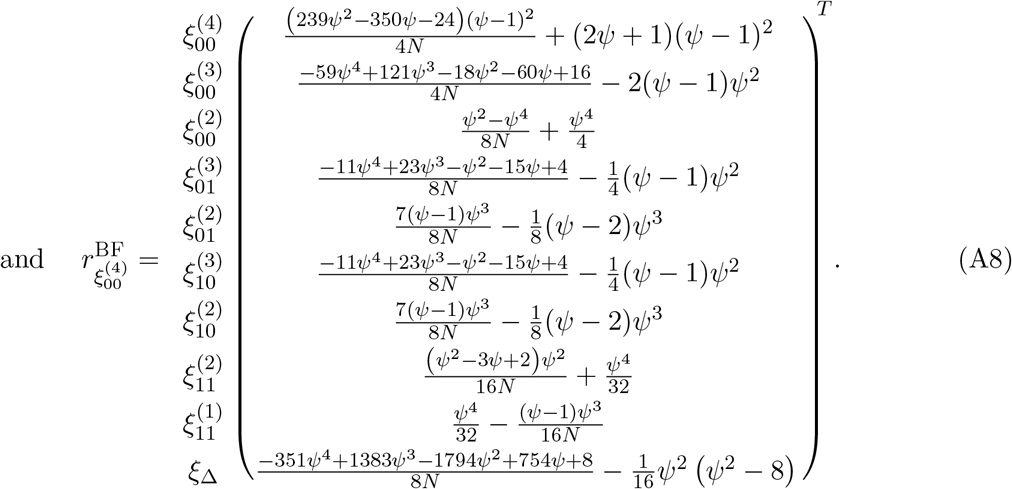

#### A.1.3 Two extremes for the sizes of big families

Degenerate case of no offspring, ***ψ*** = **0**. When a Wright-Fisher generation is taking place, denote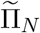 by 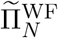. The matrix 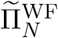 can be recovered from 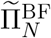 by setting *ψ* = 0 up to a term of order O(*N*^−2^). The additional O(*N*^−2^) term represents cases in which both pairs of lineages coalesce and which become negligible in the limit. Note there is a single case of non-negligible double coalescence in the O(*N*^−1^) transition 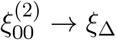. The matrix 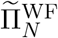 is given as

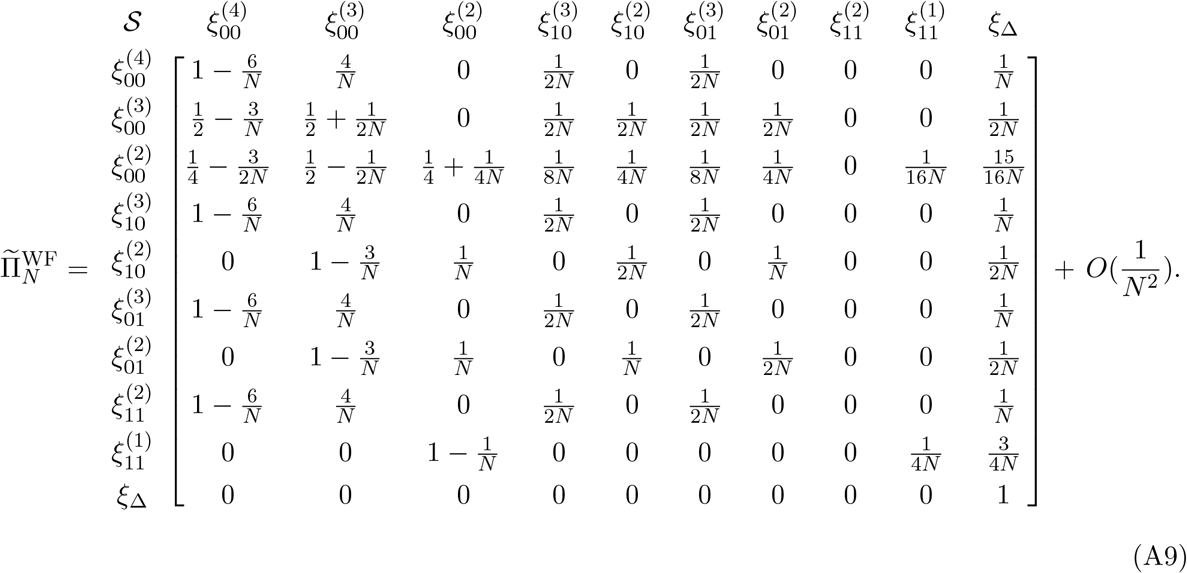

##### Offspring make up the whole population, *ψ* = 1

When *ψ* = 1, the highly reproductive pair gives birth to the entire next population. This is the case of pure Mendelian randomness within a single family with two parents. The matrix 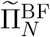 is then equal to

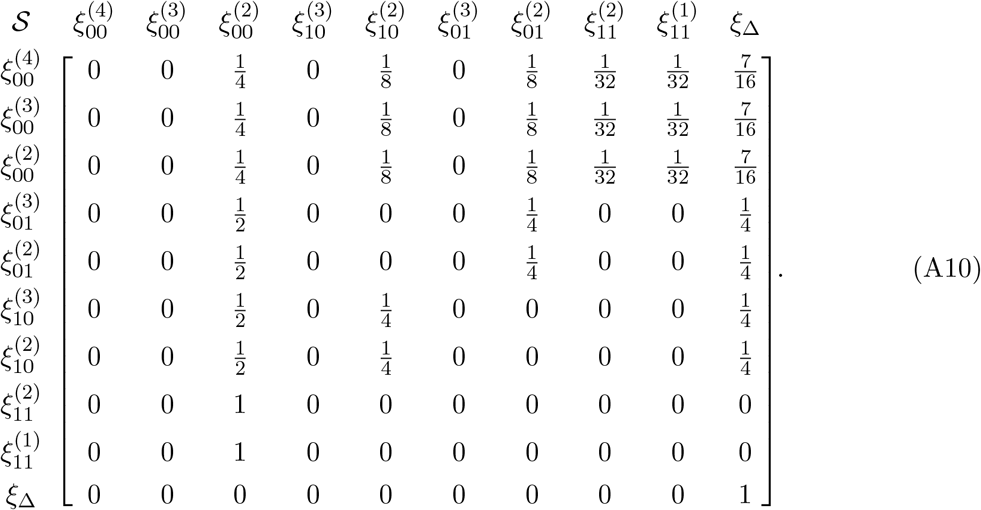

The entries of the matrix above can be explained in terms of Mendel’s laws of independent segregation and independent assortment, given that all individuals have the same two parents. For example, in the first row the initial state is 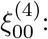 all four lineages in distinct individuals. The transition to state 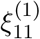, where all four lineages are in the same individual but have not coalesced, takes place with probability 1/32 and is computed as follows. Each lineage chooses a parent with probability 1/2, then chooses a gene copy within that parent also with probability 1/2. All four lineages choose the same parent (which can be either of the two) with probability 1/8. Then each pair independently chooses distinct gene copies (i.e. does not coalesce) with probability 1/2.

#### A.1.4 Limiting behavior of the joint diploid ancestral process

The joint diploid ancestral process has transition matrix

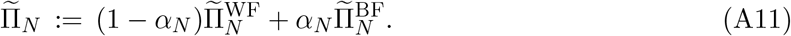

One can write

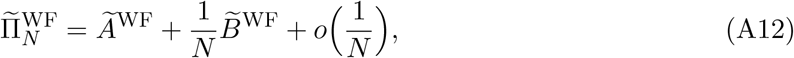

where

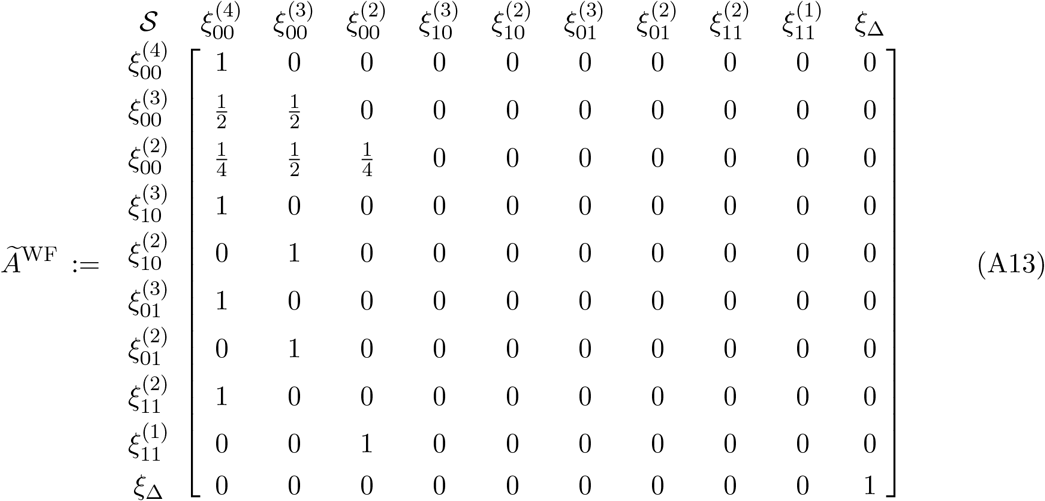

and

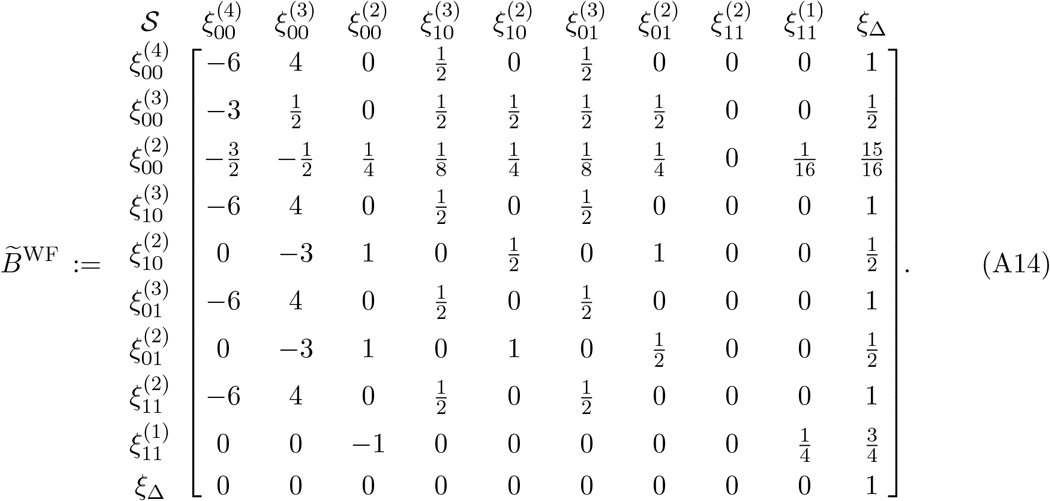

Similarly, we split

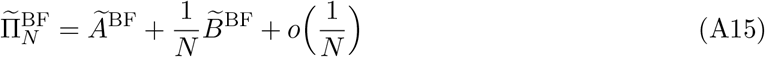

with

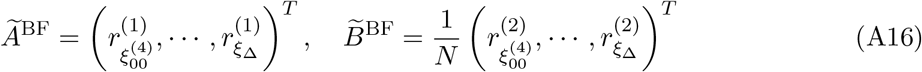

defined as 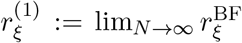 and 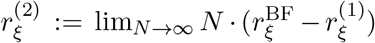 for all *ξ* ∈ *𝒮*. We provide only the rows of 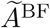 since those are the only ones we will need for the subsequent computations. In particular those are given as

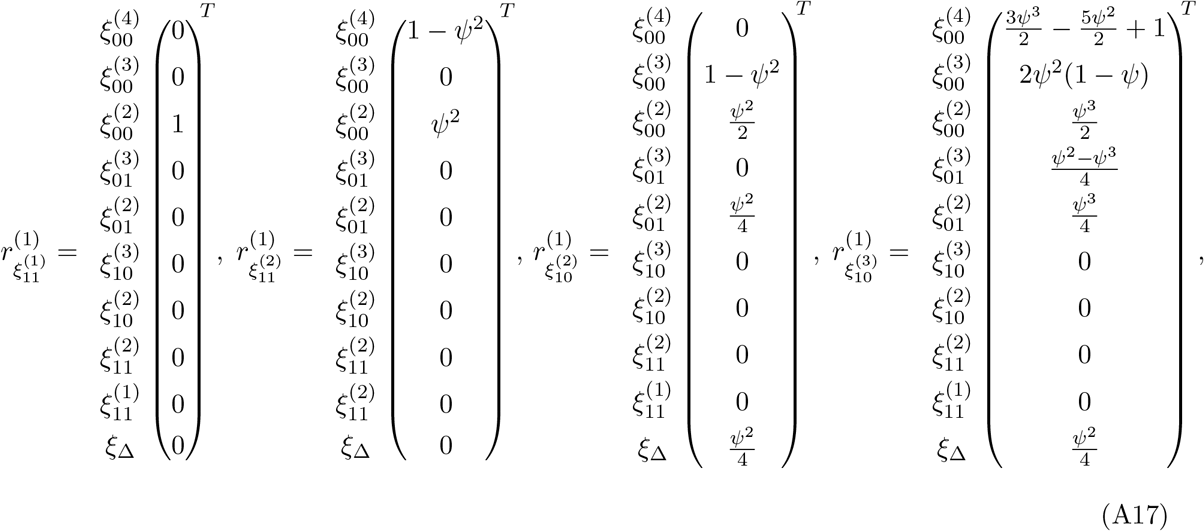

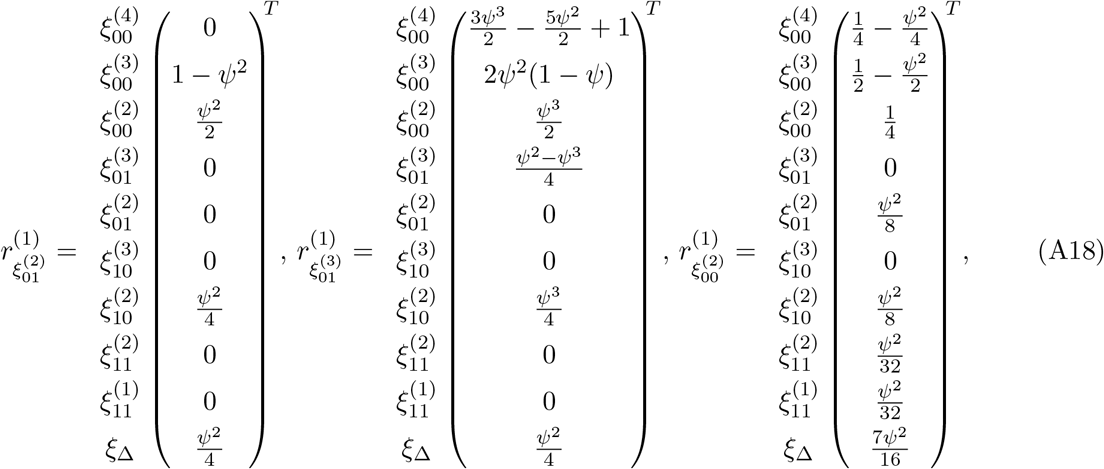

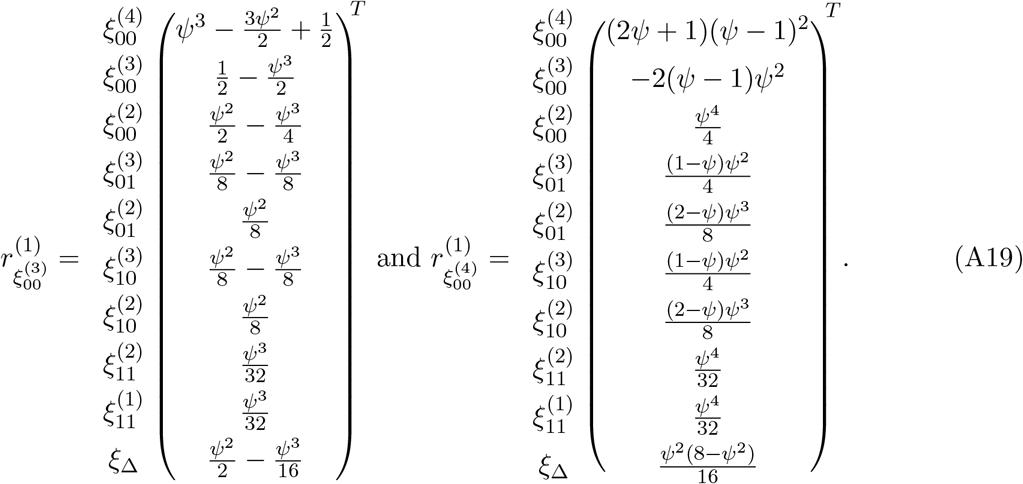

Consider now the matrices 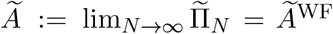 and 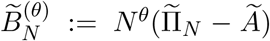. We see from (A13) that

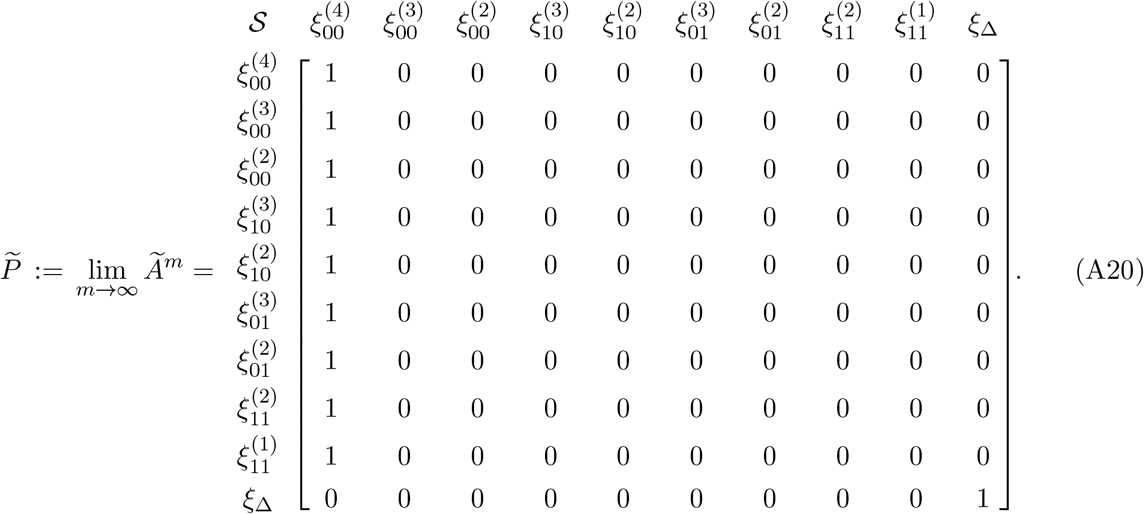

In fact, 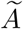 is the transition matrix of a Markov chain on 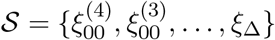 with two absorbing states 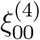and *ξ*_Δ_; furthermore, 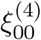 is reached from every *ξ* ∈ *𝒮 \* {*ξ*_Δ_}. Denoting 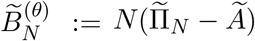 we can write 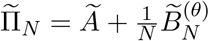, for *α*_*N*_ = *λ*/*N*^*θ*^ and *θ* ∈ (0, 1]. Also let

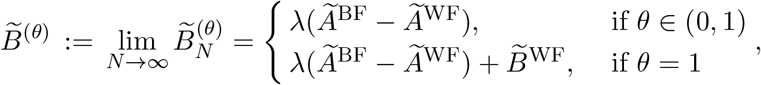

and observe that the first nine rows of

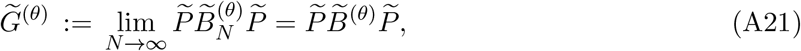

are identical and equal to

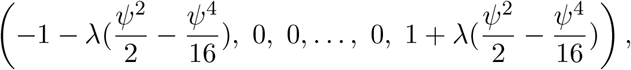

when *θ* = 1, and equal to

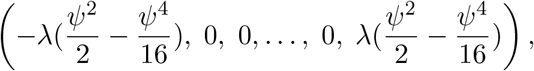

when *θ* ∈ (0, 1). The last row of *G*^(*θ*)^ is equal to (0, 0, …, 0) for all *θ* ∈ (0, 1]. Note that for computations involving 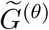, we can effectively collapse the state space 𝒮 into two points: *ξ*_Δ_ and 𝒮 *\*{*ξ*_Δ_}. Now we have all the required terms to invoke Möhle (1998a, Lemma 1) for the proof of Lemma 2. We note that sup_*N*_ 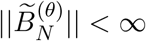, where for any matrix *K* = (*k*_*ij*_)_*i,j*_, the norm ||*K*|| := max_*i*_Σ_*j*_ |*k*_*ij*_| is used. The calculations for Lemma 2 are available at https://github.com/diamantidisdimitris/Bursts-of-coalescence in the **Lemma_2_second_moment_print** file. The computations can be reproduced using the **Lemma_2_second_moment_notebook** Mathematica notebook, version 13.1.0 (Wolfram Research, Inc., 2022).

##### Lemma 2.

Let *τ* and *τ*^*′*^ be the coalescent times of the two pairs of genes that are conditionally independent given 𝒜^(*N*,2)^. Suppose the initial distribution 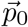 of 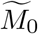 puts zero mass on state *ξ*_Δ_. Then for all 0 < *t* ≤ *t*^*′*^, as *N* → *∞*,

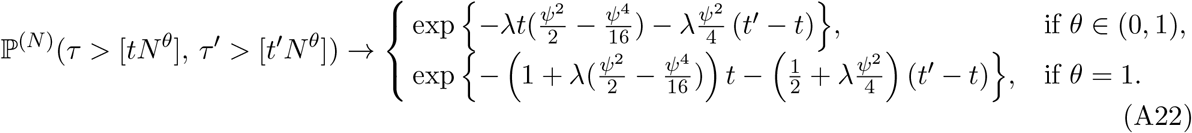

**Proof:** We first establish the result for the case *t* = *t*^*′*^, and then strengthen it to the general case *t* ≤ *t*^*′*^ by using the Markov property.

Let 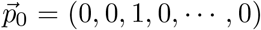 be the 1 *×* 13 row vector representing the distribution of 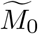, which corresponds to the initial distribution of the Markov chain. Then for each *N* ∈ *ℕ* and each generation *g* ∈ *ℤ*_≥0_,

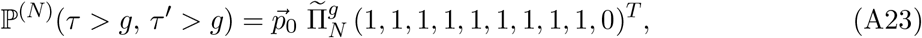

where 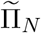 was defined in (A11). Setting *g* = [*tN*^*θ*^] in (A23), an application of Möhle (1998a, Lemma 1) gives

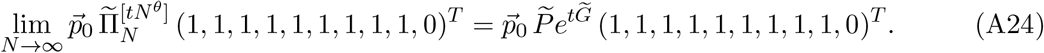

Substituting 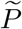 from (A20) and 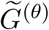 from (A21), the right hand side of (A24) is equal to

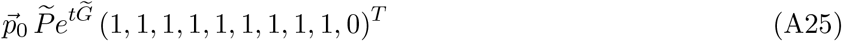

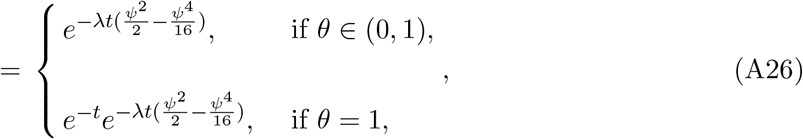

for any 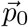 in the set 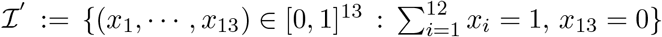. We have shown that

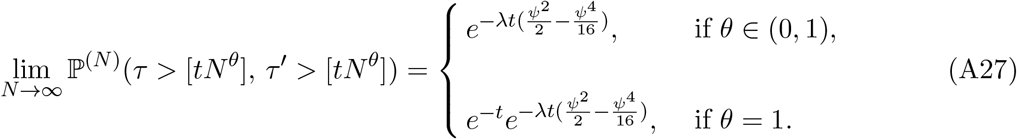

For each *N* ∈ *ℕ* and generations 0 ≤ *g* ≤ *g*^*′*^, by the Markov property,

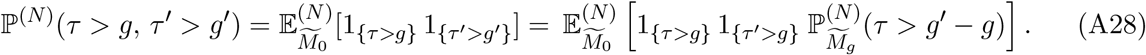

Set *g* = [*tN*^*θ*^] and *g*^*′*^ = [*t*^*′*^*N*^*θ*^]. Note that 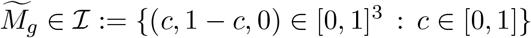when *τ* > *g*. Hence by Remark 1 and Lemma 1,

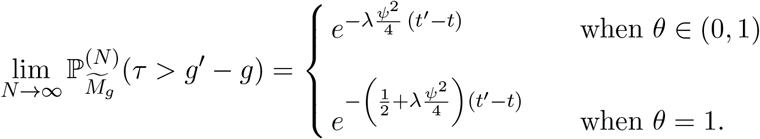

Putting this into (A28) and using (A27), we obtain the desired (A22). □

### A.2 Supporting Lemmas

The second step in the proof of Theorem 1 was the computation of (24) for *θ* = 1 and of (37) for *θ* ∈ (0, 1). An important step in those calculations was to establish (33). Lemma 3 below establishes (33). Similarly to (16), for each *g* ∈ℤ _≥0_ we let 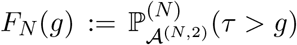 and 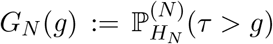.

#### Lemma 3.

Recall the definition of 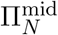 in (34). For all *g* ∈ *ℕ, ψ* ∈ (0, 1] and *N* ≥ 2 and initial distribution 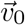 (represented as a 1 *×* 3 row vector),

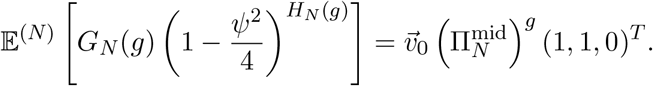

**Proof:** By the law of total expectation,

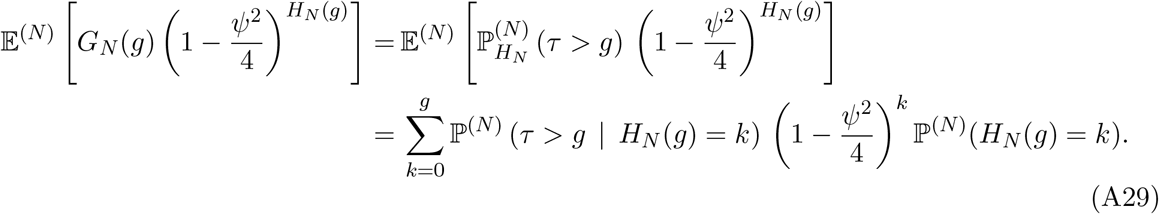

Given *g* generations, k of which being generations with big families, the total number of arrangements of those events in the pedigree is equal to 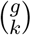. Given k, the arrangement is uniform on *W*_*k*_, the space of all possible arrangements of *k* big families within *g* generations. Denote each such arrangement by 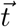. Let 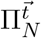 be the product of *g* terms, each term being either 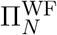 or 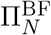. The order of the terms (from left to right) is determined by the vector 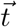 so that whenever a Wright-Fisher generation is taking place the corresponding factor in the product is 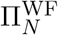 and 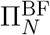 otherwise. Hence, the conditional probability in (A29) is equal to

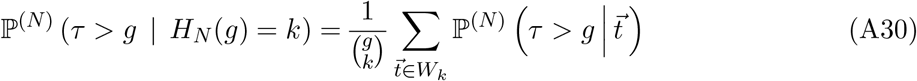

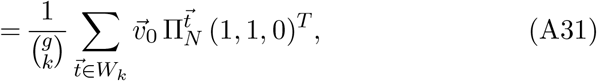

where (A30) is true by the conditional law of total probability and (A31) by the definition of conditional probability. Plugging (A31) back into (A29), since *H*_*N*_ (*g*) ∼ Bin(*g, α*_*N*_), it follows

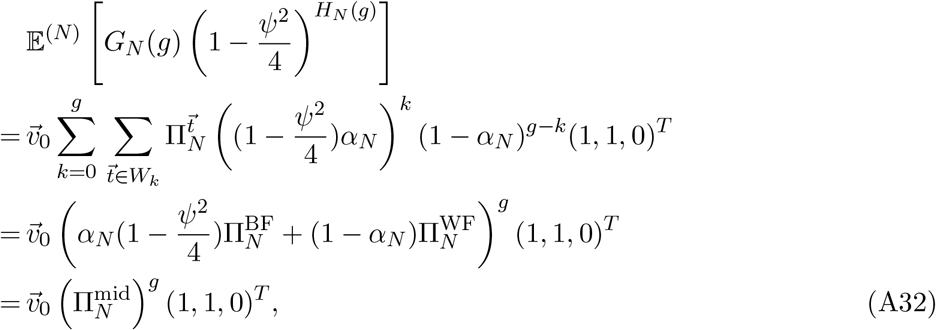

where the second last equality follows from distributing *k* terms of *α*_*N*_ and *g* − *k* terms of 1 − *α*_*N*_ into the product 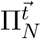. □

Finally, we compute the right hand of (33). The calculations for Lemma 4 are available at https://github.com/diamantidisdimitris/Bursts-of-coalescence in the file titled **Lemma_4_mixed_moment_print** and can be reproduced in the Mathematica notebook **Lemma_4_mixed_moment_notebook**, where Mathematica version 13.1.0 (Wolfram Research, Inc., 2022) was used.

#### Lemma 4.

Suppose the distribution 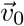 of the initial state *M*_0_ puts zero mass at the coalescent state *ξ*_2_ (recall 3-states Markov chain M in the proof of Lemma 1). For each *t* ∈ (0, *∞*), it holds that

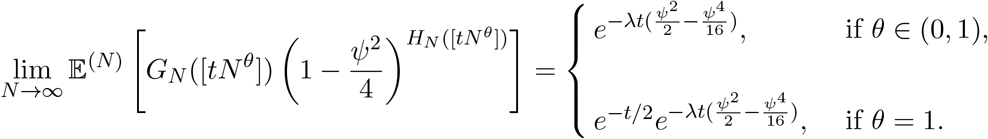

**Proof:** Let 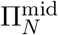 as in (34), by Lemma 3 for *g* = [*tN*^*θ*^] it follows that

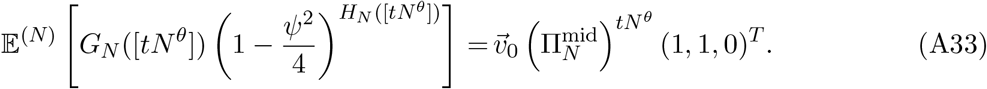

Notice that 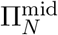 is not a stochastic matrix, consider instead

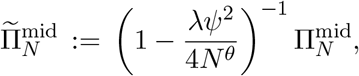

so that 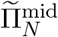 is a stochastic matrix. For any *θ* ∈ (0, 1], let

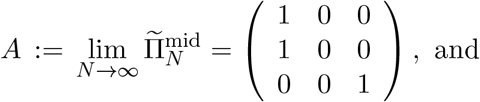

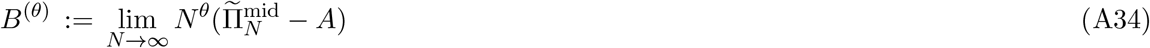

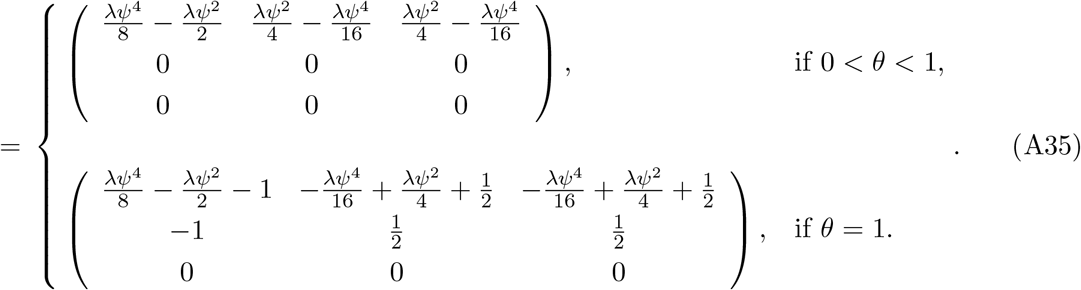

Furthermore consider the matrix *P* := lim_*k*→*∞*_ *A*^*k*^, then *P* = *A* and let *G*^(*θ*)^ := *PB*^(*θ*)^*P*. Then for *θ* ∈ (0, 1), the first two rows of *G*^(*θ*)^ are identical to

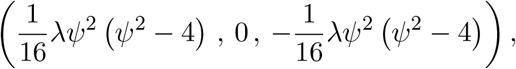

and the last one is equal to (0, 0, 0). For *θ* = 1, the first two rows of *G*^(1)^ are identical to

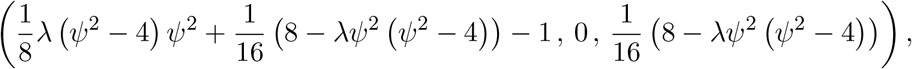

and the last one is equal to (0, 0, 0). Writing 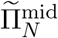 as

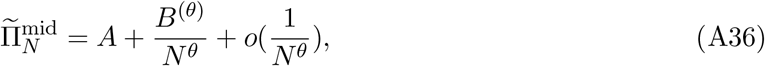

we can apply Möhle’s Lemma (Möhle, 1998a, Lemma 1) which gives

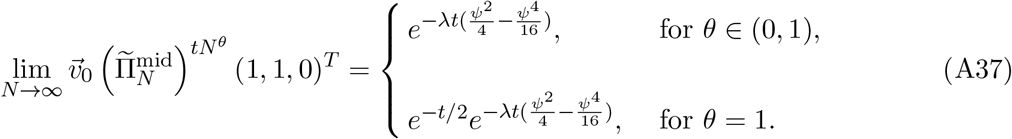

Hence (A33), through the use of (A37), is equal to

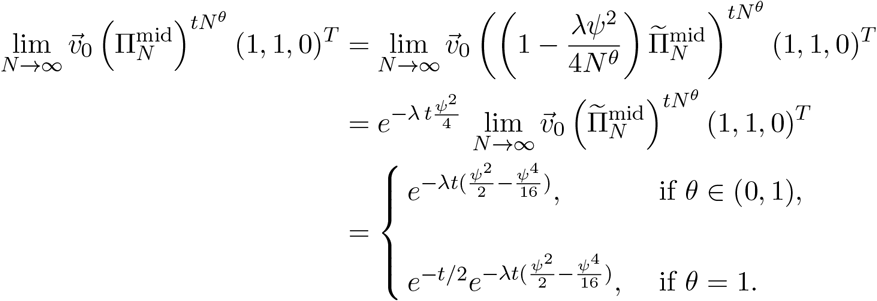

The last equality holds because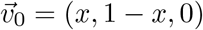 for some *x* ∈ [0, 1]. □

#### A.2.1 Proof sketch for Remark 3

Let 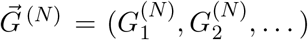 with 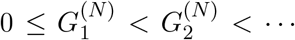 be the (random) indices of the generations when a big family occurs. By the definition of our model, we can think of generating the random pedigree in a two-step procedure: First generate 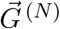 by deciding for each generation independently with probability *α*_*N*_ that a big family occurs there. Then given 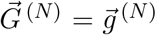 with 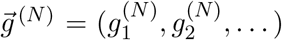 fixed, generate the pedigree between generation *g* and *g* + 1 by assigning offspring to parents according to the “generation with big family” model if 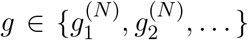 and according to the diploid Wright-Fisher model otherwise, independently for different *g*’s.

Thus, 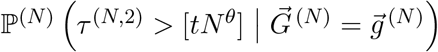 can be computed by using a time-inhomogeneous Markov chain with the three states *ξ*_0_, *ξ*_1_ and *ξ*_2_ from Lemma 1, as in (12),

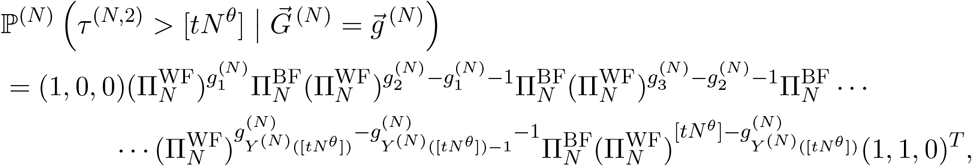

where 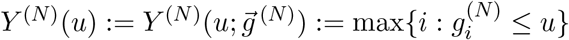.

Now 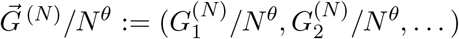 converges in distribution to (*J*_1_, *J*_2_, …) where 0 < *J*_1_ < *J*_2_ < *· · ·* are the jump times of a Poisson process (*Y* (*t*))_*t*≥0_ with rate *λ* (and *Y* (*t*) = max{*i* : J_*i*_ ≤ *t*}).

We see from (8) that

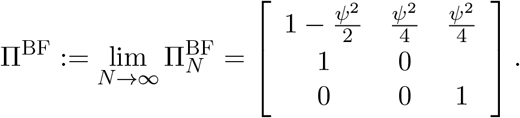

Furthermore, for *θ* = 1 we obtain from (7) that for any *t* > 0

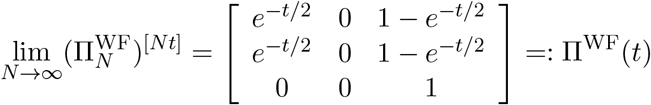

whereas for *θ* ∈ (0, 1)

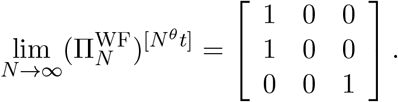

This gives for *θ* = 1

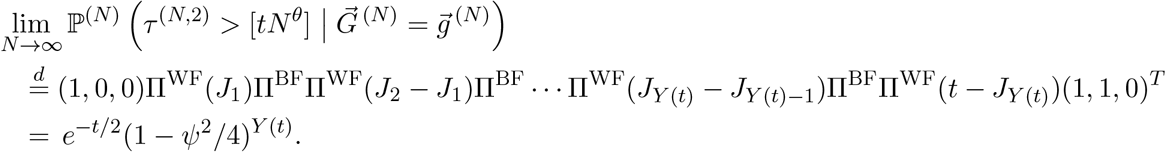

The argument for *θ* ∈ (0, 1) is analogous. □

In fact, the argument above suggests that the model and the main result can be extended to a scenario where the limiting process (*Y* (*t*)) is not necessarily a time-homogeneous Poisson process as long as the construction of the pedigree conditional on the sequence of times 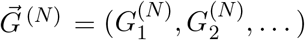 is unchanged.

### A.3 List of Patterns and Mendelian randomness tables

#### A.3.1 List of patterns

For the Tables A1-A3, let *x, y, z, w* ∈ [*N*] represent distinct individuals. Circles represent the parent(s) of these individuals. For any individual, two arrows pointing to the same circle indicate that the individual results from selfing. If two individuals each have one arrow pointing to the same circle, it means that the individuals have exactly one parent in common. The third column in each of the Tables A1-A3 contains the probabilities of the patterns in a Wright-Fisher generation, while the last column represents a generation with a big family. We only keep track of terms of order *O*(1) and *O*(*N*^−1^).

For example, the probabilities of the patterns *A*_11_ and *A*_12_ are explained as follows: In the offspring generation, an individual originates from the highly reproductive pair with probability *ψ*, signifying two distinct parents. Conversely, the same individual is *not* among the offspring of the highly reproductive pair with a probability of 1 − *ψ*. In this scenario, the individual comes from selfing with a probability of 1/*N*, and from two distinct parents with probability 1 − 1/*N*. Finally, the probability of pattern *A*_22_ is explained as follows: Each individual in the offspring generation originates from the highly reproductive pair with probability *ψ*. By independence, both individuals are offspring of the highly reproductive couple with probability *ψ*^2^. Note that two individuals can share both their parents in various ways. However, in all other scenarios, the probability of such an event is of the order *O*(*N*^−2^).

**Table A1:**
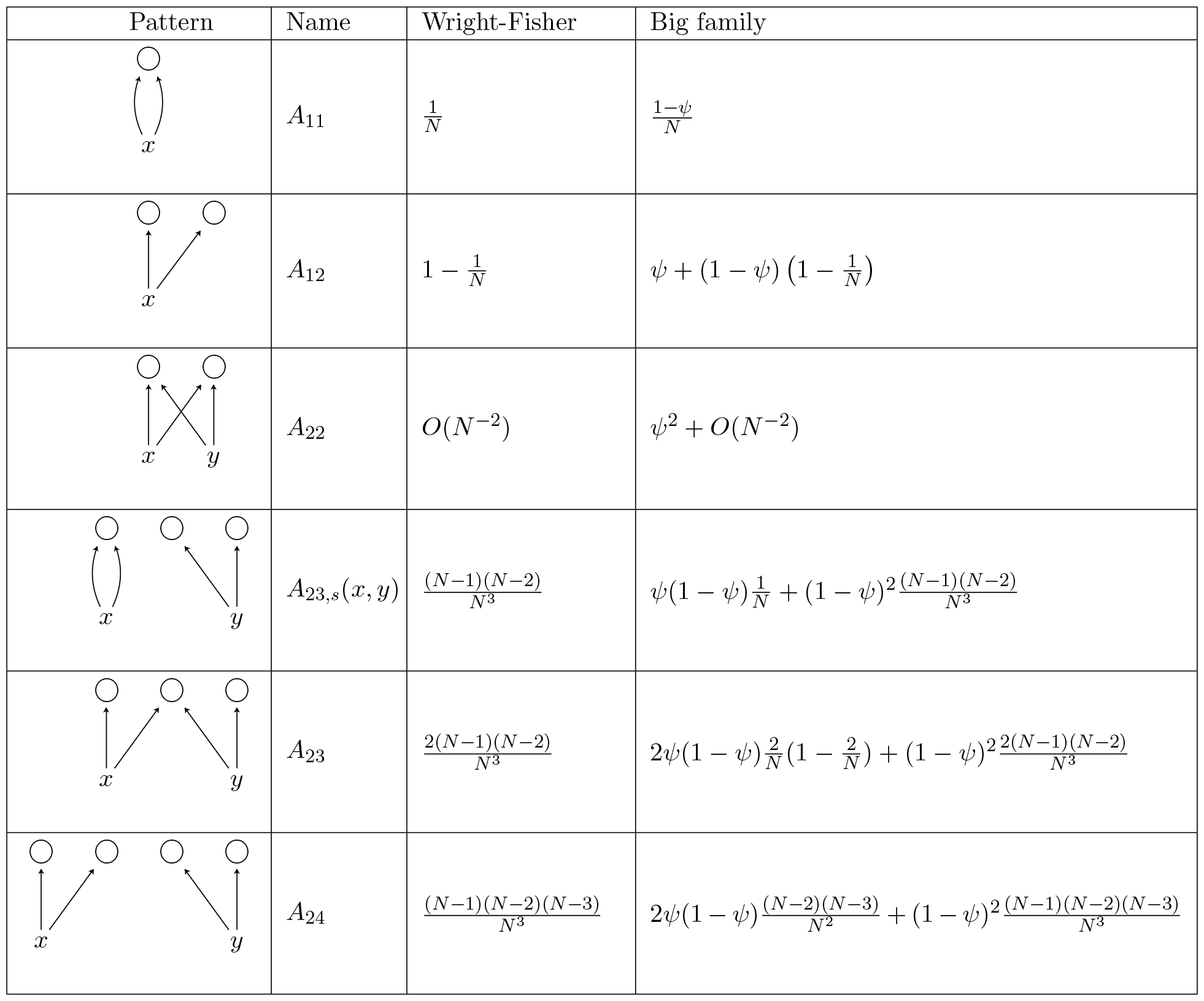
List of patterns when the sampled genes are distributed among one or two individuals.

| Pattern | Name | Wright-Fisher | Big family |
| --- | --- | --- | --- |
| | $A_{11}$ | $\frac{1}{N}$ | $\frac{1-\psi}{N}$ |
| | $A_{12}$ | $1 - \frac{1}{N}$ | $\psi + (1 - \psi) \left(1 - \frac{1}{N}\right)$ |
| | $A_{22}$ | $O(N^{-2})$ | $\psi^2 + O(N^{-2})$ |
| | $A_{23,s}(x, y)$ | $\frac{(N-1)(N-2)}{N^3}$ | $\psi(1 - \psi) \frac{1}{N} + (1 - \psi)^2 \frac{(N-1)(N-2)}{N^3}$ |
| | $A_{23}$ | $\frac{2(N-1)(N-2)}{N^3}$ | $2\psi(1 - \psi) \frac{2}{N} \left(1 - \frac{2}{N}\right) + (1 - \psi)^2 \frac{2(N-1)(N-2)}{N^3}$ |
| | $A_{24}$ | $\frac{(N-1)(N-2)(N-3)}{N^3}$ | $2\psi(1 - \psi) \frac{(N-2)(N-3)}{N^2} + (1 - \psi)^2 \frac{(N-1)(N-2)(N-3)}{N^3}$ |

**Table A2:**
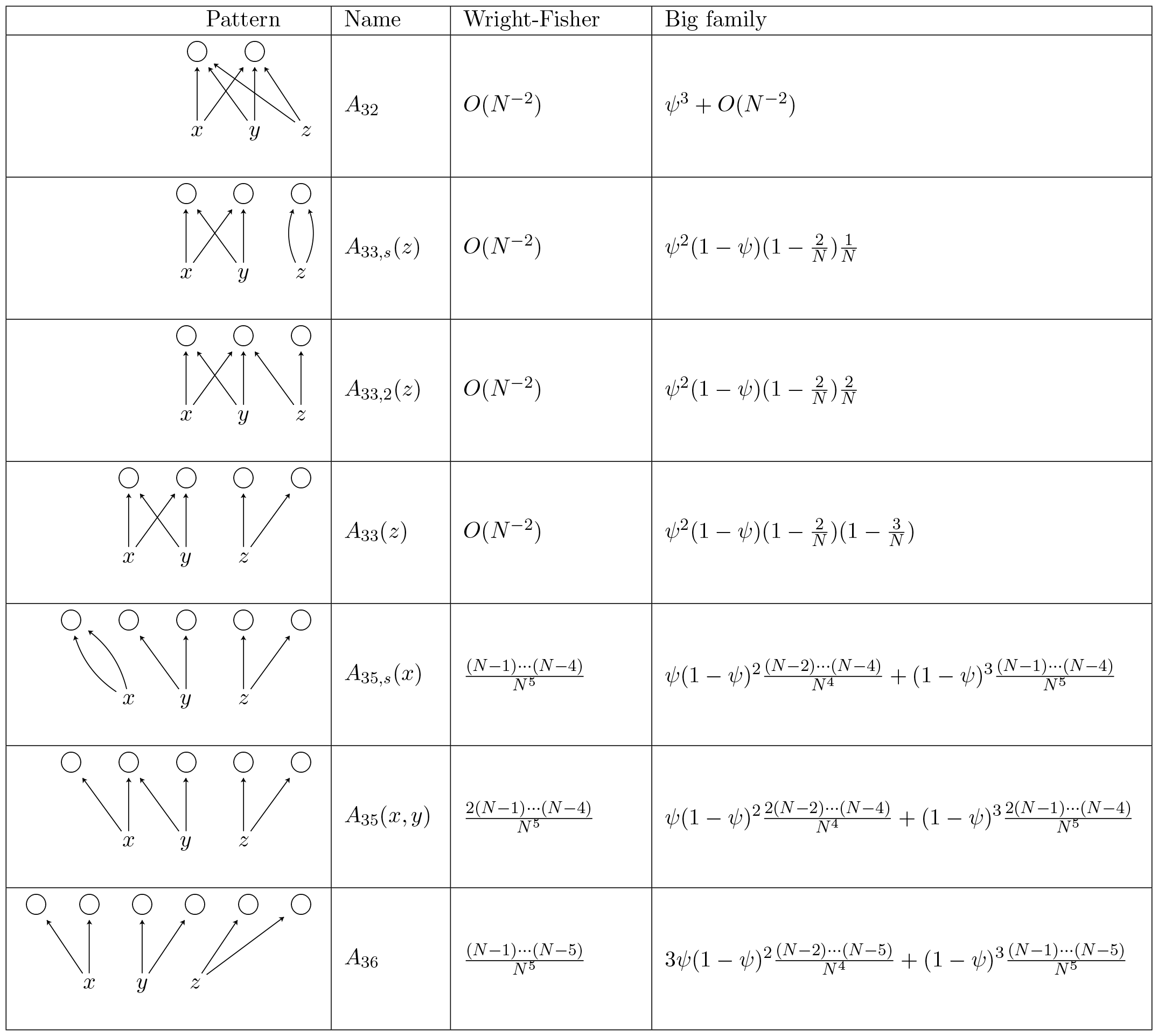
List of patterns when the sampled genes are distributed among three individuals.

| Pattern | Name | Wright-Fisher | Big family |
| --- | --- | --- | --- |
| | $A_{32}$ | $O(N^{-2})$ | $\psi^3 + O(N^{-2})$ |
| | $A_{33,s}(z)$ | $O(N^{-2})$ | $\psi^2(1-\psi)(1-\frac{2}{N})\frac{1}{N}$ |
| | $A_{33,2}(z)$ | $O(N^{-2})$ | $\psi^2(1-\psi)(1-\frac{2}{N})\frac{2}{N}$ |
| | $A_{33}(z)$ | $O(N^{-2})$ | $\psi^2(1-\psi)(1-\frac{2}{N})(1-\frac{3}{N})$ |
| | $A_{35,s}(x)$ | $\frac{(N-1)\cdots(N-4)}{N^5}$ | $\psi(1-\psi)^2\frac{(N-2)\cdots(N-4)}{N^4} + (1-\psi)^3\frac{(N-1)\cdots(N-4)}{N^5}$ |
| | $A_{35}(x, y)$ | $\frac{2(N-1)\cdots(N-4)}{N^5}$ | $\psi(1-\psi)^2\frac{2(N-2)\cdots(N-4)}{N^4} + (1-\psi)^3\frac{2(N-1)\cdots(N-4)}{N^5}$ |
| | $A_{36}$ | $\frac{(N-1)\cdots(N-5)}{N^5}$ | $3\psi(1-\psi)^2\frac{(N-2)\cdots(N-5)}{N^4} + (1-\psi)^3\frac{(N-1)\cdots(N-5)}{N^5}$ |

**Table A3:**
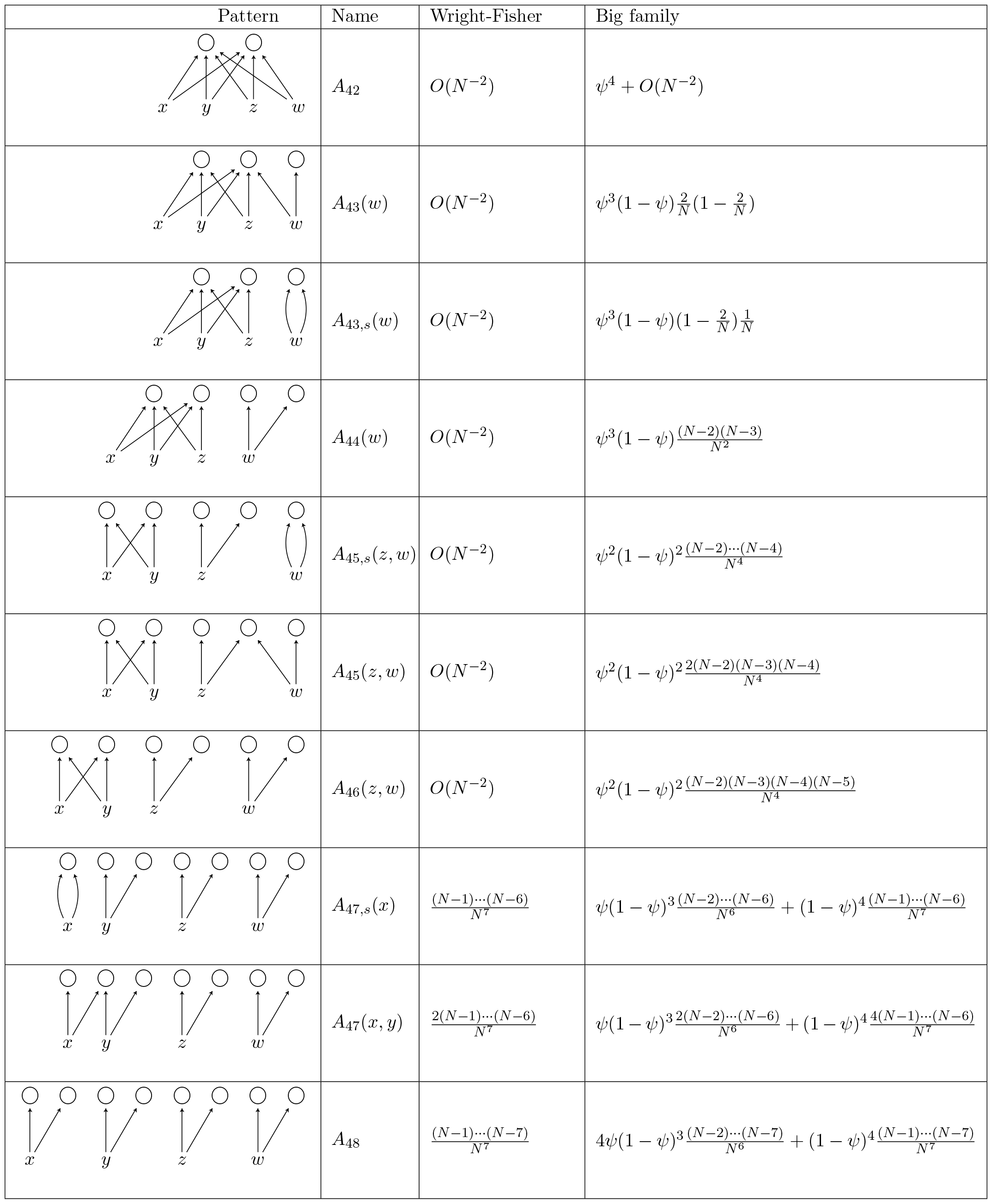
List of patterns when the sampled genes are distributed among four individuals.

| Pattern | Name | Wright-Fisher | Big family |
| --- | --- | --- | --- |
| | $A_{42}$ | $O(N^{-2})$ | $\psi^4 + O(N^{-2})$ |
| | $A_{43}(w)$ | $O(N^{-2})$ | $\psi^3(1 - \psi) \frac{2}{N} (1 - \frac{2}{N})$ |
| | $A_{43,s}(w)$ | $O(N^{-2})$ | $\psi^3(1 - \psi)(1 - \frac{2}{N}) \frac{1}{N}$ |
| | $A_{44}(w)$ | $O(N^{-2})$ | $\psi^3(1 - \psi) \frac{(N-2)(N-3)}{N^2}$ |
| | $A_{45,s}(z, w)$ | $O(N^{-2})$ | $\psi^2(1 - \psi)^2 \frac{(N-2) \cdots (N-4)}{N^4}$ |
| | $A_{45}(z, w)$ | $O(N^{-2})$ | $\psi^2(1 - \psi)^2 \frac{2(N-2)(N-3)(N-4)}{N^4}$ |
| | $A_{46}(z, w)$ | $O(N^{-2})$ | $\psi^2(1 - \psi)^2 \frac{(N-2)(N-3)(N-4)(N-5)}{N^4}$ |
| | $A_{47,s}(x)$ | $\frac{(N-1) \cdots (N-6)}{N^7}$ | $\psi(1 - \psi)^3 \frac{(N-2) \cdots (N-6)}{N^6} + (1 - \psi)^4 \frac{(N-1) \cdots (N-6)}{N^7}$ |
| | $A_{47}(x, y)$ | $\frac{2(N-1) \cdots (N-6)}{N^7}$ | $\psi(1 - \psi)^3 \frac{2(N-2) \cdots (N-6)}{N^6} + (1 - \psi)^4 \frac{4(N-1) \cdots (N-6)}{N^7}$ |
| | $A_{48}$ | $\frac{(N-1) \cdots (N-7)}{N^7}$ | $4\psi(1 - \psi)^3 \frac{(N-2) \cdots (N-7)}{N^6} + (1 - \psi)^4 \frac{(N-1) \cdots (N-7)}{N^7}$ |

#### A.3.2 Mendelian randomness tables

Tables A4-A10 below, contain the transition probabilities for every pair of states (*ξ, η*) ∈ *𝒮*^2^, conditional on the patterns from Tables A1-A3. The derivation of the entries of Tables A4-A10 follow similarly to the entries of (A10) in subsection A.1.3. For example, in Table A10, the last row of the column labeled *A*_47_ is explained as follows. Lineages in distinct individuals sharing exactly one parent have descended from different parents with probability 3/4. On the other hand, state 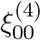 consists of two distinct individuals, hence the pattern *A*_47_ is used in 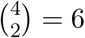 ways. Finally we note that in Tables A4-A10, the fractional quantities stand for events determined purely by Mendelian randomness (conditional on each pattern) whereas the integer quantities refer to combinatorial terms corresponding to the appropriate use of the patterns.

**Table A4:** Conditional probabilities given the pattern when the initial state is 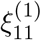.

| $(\xi_{11}^{(1)}, \eta)$ | $A_{11}$ | $A_{12}$ |
| --- | --- | --- |
| $(\xi_{11}^{(1)}, \xi_{11}^{(1)})$ | $\frac{1}{4}$ | 0 |
| $(\xi_{11}^{(1)}, \xi_{11}^{(2)})$ | 0 | 0 |
| $(\xi_{11}^{(1)}, \xi_{10}^{(2)})$ | 0 | 0 |
| $(\xi_{11}^{(1)}, \xi_{10}^{(3)})$ | 0 | 0 |
| $(\xi_{11}^{(1)}, \xi_{01}^{(2)})$ | 0 | 0 |
| $(\xi_{11}^{(1)}, \xi_{01}^{(3)})$ | 0 | 0 |
| $(\xi_{11}^{(1)}, \xi_{00}^{(2)})$ | 0 | $\frac{1}{1}$ |
| $(\xi_{11}^{(1)}, \xi_{00}^{(3)})$ | 0 | 0 |
| $(\xi_{11}^{(1)}, \xi_{00}^{(4)})$ | 0 | 0 |

**Table A5:**
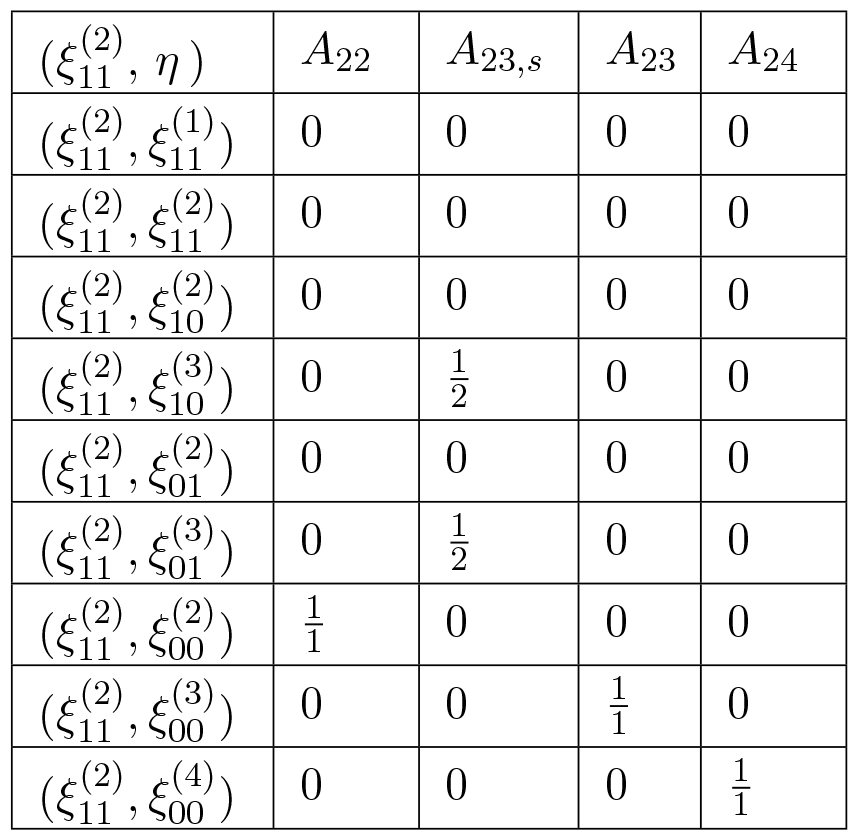
Conditional probabilities given the pattern when the initial state is 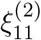.

**Table A6:** Conditional probabilities given the pattern when the initial state is 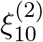.

| $(\xi_{10}^{(2)}, \eta)$ | $A_{22}$ | $A_{23,s}$ | $A_{23}$ | $A_{24}$ |
| --- | --- | --- | --- | --- |
| $(\xi_{10}^{(2)}, \xi_{11}^{(1)})$ | 0 | 0 | 0 | 0 |
| $(\xi_{10}^{(2)}, \xi_{11}^{(2)})$ | 0 | 0 | 0 | 0 |
| $(\xi_{10}^{(2)}, \xi_{10}^{(2)})$ | 0 | $\frac{1}{2}$ | 0 | 0 |
| $(\xi_{10}^{(2)}, \xi_{10}^{(3)})$ | 0 | 0 | 0 | 0 |
| $(\xi_{10}^{(2)}, \xi_{01}^{(2)})$ | $\frac{1}{4}$ | 0 | $\frac{1}{4}$ | 0 |
| $(\xi_{10}^{(2)}, \xi_{01}^{(3)})$ | 0 | 0 | 0 | 0 |
| $(\xi_{10}^{(2)}, \xi_{00}^{(2)})$ | $\frac{1}{2}$ | 0 | $\frac{1}{4}$ | 0 |
| $(\xi_{10}^{(2)}, \xi_{00}^{(3)})$ | 0 | $1 \cdot \frac{1}{1}$ | $\frac{1}{2}$ | $\frac{1}{1}$ |
| $(\xi_{10}^{(2)}, \xi_{00}^{(4)})$ | 0 | 0 | 0 | 0 |

**Table A7:** Conditional probabilities given the pattern when the initial state is 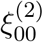.

| $(\xi_{00}^{(2)}, \eta)$ | $A_{22}$ | $A_{23,s}$ | $A_{23}$ | $A_{24}$ |
| --- | --- | --- | --- | --- |
| $(\xi_{00}^{(2)}, \xi_{11}^{(1)})$ | $\frac{1}{32}$ | 0 | $\frac{1}{64}$ | 0 |
| $(\xi_{00}^{(2)}, \xi_{11}^{(2)})$ | $\frac{1}{32}$ | 0 | 0 | 0 |
| $(\xi_{00}^{(2)}, \xi_{10}^{(2)})$ | $\frac{1}{8}$ | 0 | $\frac{1}{16}$ | 0 |
| $(\xi_{00}^{(2)}, \xi_{10}^{(3)})$ | 0 | 0 | $\frac{1}{32}$ | 0 |
| $(\xi_{00}^{(2)}, \xi_{01}^{(2)})$ | $\frac{1}{8}$ | 0 | $\frac{1}{16}$ | 0 |
| $(\xi_{00}^{(2)}, \xi_{01}^{(3)})$ | 0 | 0 | $\frac{1}{32}$ | 0 |
| $(\xi_{00}^{(2)}, \xi_{00}^{(2)})$ | $\frac{1}{4}$ | $2 \cdot \frac{1}{2}$ | $1 \cdot \frac{3}{16}$ | $\frac{1}{4}$ |
| $(\xi_{00}^{(2)}, \xi_{00}^{(3)})$ | 0 | $2 \cdot \frac{1}{2}$ | $1 \cdot \frac{6}{16}$ | $\frac{1}{2}$ |
| $(\xi_{00}^{(2)}, \xi_{00}^{(4)})$ | 0 | 0 | 0 | $\frac{1}{4}$ |

**Table A8:** Conditional probabilities given the pattern when the initial state is 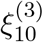.

| $(\xi_{10}^{(3)}, \eta)$ | $A_{32}$ | $A_{33,s}$ | $A_{33,2}$ | $A_{33}$ | $A_{35,s}$ | $A_{35}$ | $A_{36}$ |
| --- | --- | --- | --- | --- | --- | --- | --- |
| $(\xi_{10}^{(3)}, \xi_{11}^{(1)})$ | 0 | 0 | 0 | 0 | 0 | 0 | 0 |
| $(\xi_{10}^{(3)}, \xi_{11}^{(2)})$ | 0 | $\frac{1}{4}$ | 0 | 0 | 0 | 0 | 0 |
| $(\xi_{10}^{(3)}, \xi_{10}^{(2)})$ | 0 | 0 | 0 | 0 | 0 | 0 | 0 |
| $(\xi_{10}^{(3)}, \xi_{10}^{(3)})$ | 0 | $\frac{1}{4}$ | 0 | 0 | $1 \cdot \frac{1}{2}$ | 0 | 0 |
| $(\xi_{10}^{(3)}, \xi_{01}^{(2)})$ | $\frac{1}{4}$ | 0 | $1 \cdot \frac{1}{8}$ | 0 | 0 | 0 | 0 |
| $(\xi_{10}^{(3)}, \xi_{01}^{(3)})$ | 0 | 0 | $3 \cdot \frac{1}{8}$ | $1 \cdot \frac{1}{4}$ | 0 | $1 \cdot \frac{1}{8}$ | 0 |
| $(\xi_{10}^{(3)}, \xi_{00}^{(2)})$ | $\frac{1}{2}$ | 0 | $2 \cdot \frac{1}{4}$ | 0 | 0 | 0 | 0 |
| $(\xi_{10}^{(3)}, \xi_{00}^{(3)})$ | 0 | $2 \cdot 1$ | $3 \cdot \frac{1}{2}$ | $2 \cdot 1$ | 0 | $2 \cdot \frac{1}{2}$ | 0 |
| $(\xi_{10}^{(3)}, \xi_{00}^{(4)})$ | 0 | 0 | 0 | $1 \cdot \frac{1}{2}$ | $2 \cdot \frac{1}{1}$ | $1 \cdot \frac{3}{4} + 2 \cdot \frac{1}{2}$ | $\frac{1}{1}$ |

**Table A9:** Conditional probabilities given the pattern when the initial state is 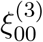.

| $(\xi_{00}^{(3)}, \eta)$ | $A_{32}$ | $A_{33,s}$ | $A_{33,2}$ | $A_{33}$ | $A_{35,s}$ | $A_{35}$ | $A_{36}$ |
| --- | --- | --- | --- | --- | --- | --- | --- |
| $(\xi_{00}^{(3)}, \xi_{11}^{(1)})$ | $1 \cdot \frac{1}{32}$ | 0 | $3 \cdot \frac{1}{64}$ | 0 | 0 | 0 | 0 |
| $(\xi_{00}^{(3)}, \xi_{11}^{(2)})$ | $1 \cdot \frac{1}{32}$ | 0 | $2 \cdot \frac{1}{64}$ | 0 | 0 | 0 | 0 |
| $(\xi_{00}^{(3)}, \xi_{10}^{(2)})$ | $1 \cdot \frac{1}{8}$ | $1 \cdot \frac{1}{8}$ | $2 \cdot \frac{1}{8} + 1 \cdot \frac{1}{16}$ | $1 \cdot \frac{1}{8}$ | 0 | $1 \cdot \frac{1}{8}$ | 0 |
| $(\xi_{00}^{(3)}, \xi_{10}^{(3)})$ | 0 | $1 \cdot \frac{1}{8}$ | $1 \cdot \frac{1}{32} + 1 \cdot \frac{1}{16}$ | $1 \cdot \frac{1}{8}$ | 0 | $1 \cdot \frac{1}{8}$ | 0 |
| $(\xi_{00}^{(3)}, \xi_{01}^{(2)})$ | $1 \cdot \frac{1}{8}$ | $1 \cdot \frac{1}{8}$ | $2 \cdot \frac{1}{16} + 1 \cdot \frac{1}{8}$ | $1 \cdot \frac{1}{8}$ | 0 | $1 \cdot \frac{1}{8}$ | 0 |
| $(\xi_{00}^{(3)}, \xi_{01}^{(3)})$ | 0 | $1 \cdot \frac{1}{8}$ | $1 \cdot \frac{1}{32} + 1 \cdot \frac{1}{16}$ | $1 \cdot \frac{1}{8}$ | 0 | $1 \cdot \frac{1}{8}$ | 0 |
| $(\xi_{00}^{(3)}, \xi_{00}^{(2)})$ | $1 \cdot \frac{1}{4}$ | $1 \cdot \frac{1}{2}$ | $1 \cdot \frac{3}{8} + 2 \cdot \frac{1}{8}$ | $1 \cdot \frac{1}{2}$ | 0 | 0 | 0 |
| $(\xi_{00}^{(3)}, \xi_{00}^{(3)})$ | 0 | $3 \cdot \frac{1}{2}$ | $1 \cdot \frac{3}{8} + 2 \cdot \frac{1}{4}$ | $3 \cdot \frac{1}{2}$ | $1 \cdot \frac{1}{1} + 2 \cdot \frac{1}{2}$ | $1 \cdot \frac{1}{2} + 2 \cdot \frac{1}{2}$ | $1 \cdot \frac{1}{2}$ |
| $(\xi_{00}^{(3)}, \xi_{00}^{(4)})$ | 0 | 0 | 0 | 0 | $2 \cdot \frac{1}{2}$ | $1 \cdot \frac{3}{8} + 2 \cdot \frac{1}{4}$ | $1 \cdot \frac{1}{2}$ |

**Table A10:**
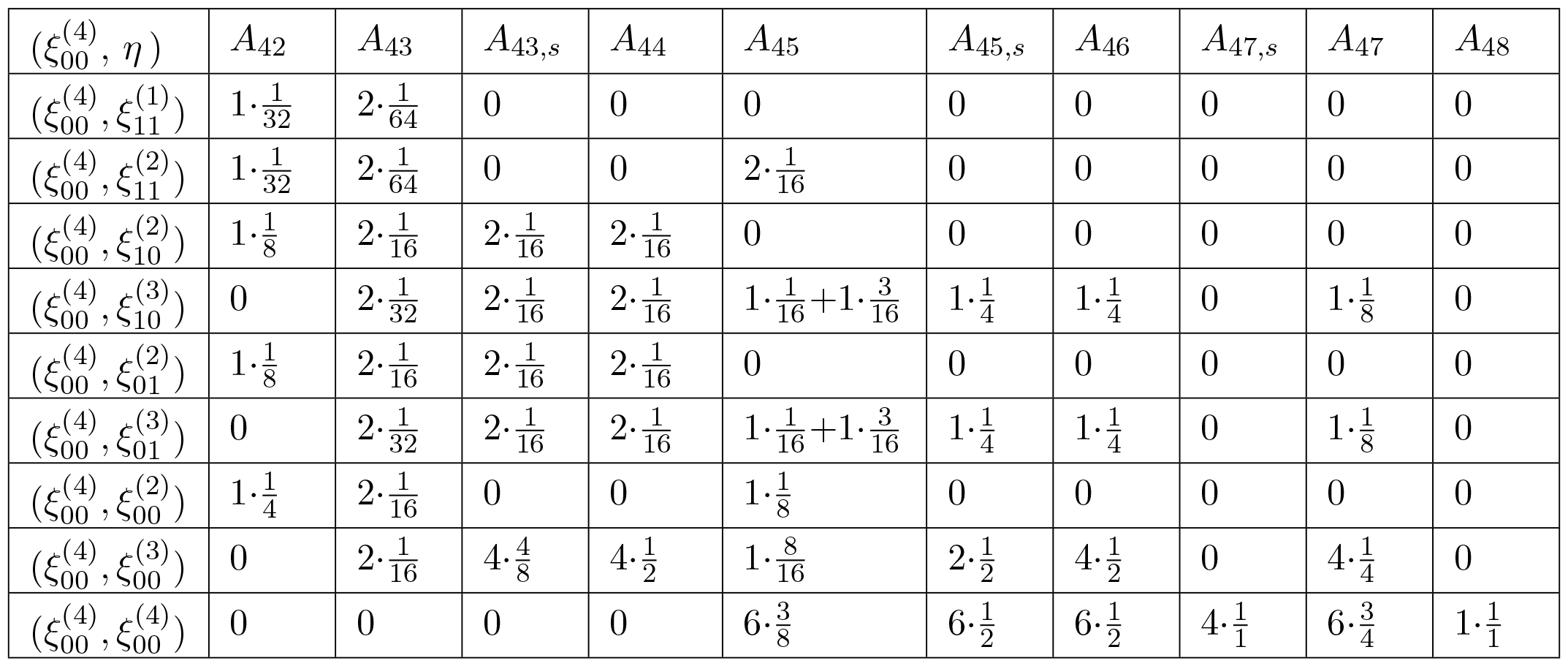
Conditional probabilities given the pattern when the initial state is 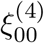.

| $(\xi_{00}^{(4)}, \eta)$ | $A_{42}$ | $A_{43}$ | $A_{43,s}$ | $A_{44}$ | $A_{45}$ | $A_{45,s}$ | $A_{46}$ | $A_{47,s}$ | $A_{47}$ | $A_{48}$ |
| --- | --- | --- | --- | --- | --- | --- | --- | --- | --- | --- |
| $(\xi_{00}^{(4)}, \xi_{11}^{(1)})$ | $1 \cdot \frac{1}{32}$ | $2 \cdot \frac{1}{64}$ | 0 | 0 | 0 | 0 | 0 | 0 | 0 | 0 |
| $(\xi_{00}^{(4)}, \xi_{11}^{(2)})$ | $1 \cdot \frac{1}{32}$ | $2 \cdot \frac{1}{64}$ | 0 | 0 | $2 \cdot \frac{1}{16}$ | 0 | 0 | 0 | 0 | 0 |
| $(\xi_{00}^{(4)}, \xi_{10}^{(2)})$ | $1 \cdot \frac{1}{8}$ | $2 \cdot \frac{1}{16}$ | $2 \cdot \frac{1}{16}$ | $2 \cdot \frac{1}{16}$ | 0 | 0 | 0 | 0 | 0 | 0 |
| $(\xi_{00}^{(4)}, \xi_{10}^{(3)})$ | 0 | $2 \cdot \frac{1}{32}$ | $2 \cdot \frac{1}{16}$ | $2 \cdot \frac{1}{16}$ | $1 \cdot \frac{1}{16} + 1 \cdot \frac{3}{16}$ | $1 \cdot \frac{1}{4}$ | $1 \cdot \frac{1}{4}$ | 0 | $1 \cdot \frac{1}{8}$ | 0 |
| $(\xi_{00}^{(4)}, \xi_{01}^{(2)})$ | $1 \cdot \frac{1}{8}$ | $2 \cdot \frac{1}{16}$ | $2 \cdot \frac{1}{16}$ | $2 \cdot \frac{1}{16}$ | 0 | 0 | 0 | 0 | 0 | 0 |
| $(\xi_{00}^{(4)}, \xi_{01}^{(3)})$ | 0 | $2 \cdot \frac{1}{32}$ | $2 \cdot \frac{1}{16}$ | $2 \cdot \frac{1}{16}$ | $1 \cdot \frac{1}{16} + 1 \cdot \frac{3}{16}$ | $1 \cdot \frac{1}{4}$ | $1 \cdot \frac{1}{4}$ | 0 | $1 \cdot \frac{1}{8}$ | 0 |
| $(\xi_{00}^{(4)}, \xi_{00}^{(2)})$ | $1 \cdot \frac{1}{4}$ | $2 \cdot \frac{1}{16}$ | 0 | 0 | $1 \cdot \frac{1}{8}$ | 0 | 0 | 0 | 0 | 0 |
| $(\xi_{00}^{(4)}, \xi_{00}^{(3)})$ | 0 | $2 \cdot \frac{1}{16}$ | $4 \cdot \frac{4}{8}$ | $4 \cdot \frac{1}{2}$ | $1 \cdot \frac{8}{16}$ | $2 \cdot \frac{1}{2}$ | $4 \cdot \frac{1}{2}$ | 0 | $4 \cdot \frac{1}{4}$ | 0 |
| $(\xi_{00}^{(4)}, \xi_{00}^{(4)})$ | 0 | 0 | 0 | 0 | $6 \cdot \frac{3}{8}$ | $6 \cdot \frac{1}{2}$ | $6 \cdot \frac{1}{2}$ | $4 \cdot \frac{1}{1}$ | $6 \cdot \frac{3}{4}$ | $1 \cdot \frac{1}{1}$ |

### A.4 Strengthening the convergence in Theorem 1

#### Remark 4.

*The arguments used in the proof of Theorem 1 in fact allow to conclude the (stronger) statement that*

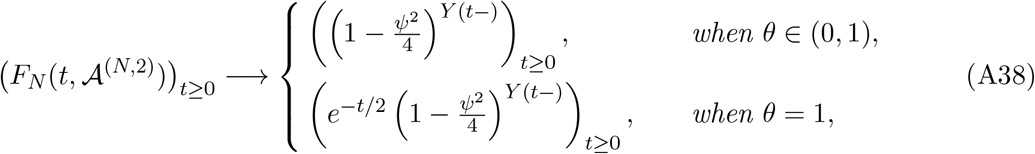

*in finite-dimensional distributions*.

*Proof*. We will only consider the case *θ* = 1 here, the case *θ* < 1 is similar.

Fix *k* ∈ *ℕ* and 0 < *t*_1_ < *t*_2_ < *· · · t*_*k*_. Using (21) for *t*_1_, …, *t*_*k*_, we have

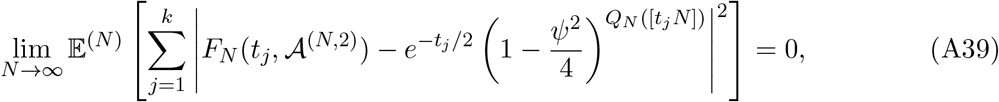

which implies as in the proof of Theorem 1 that

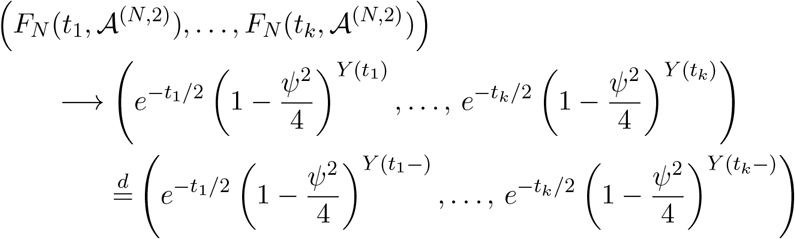

in distribution. This shows convergence of finite dimensional distributions. □

Further we conjecture that we have weak convergence in the space of left-continuous functions with right limits, equipped with the Skorohod topology, but we do not pursue this here.

